# A Novel Rapid Host Cell Entry Pathway Determines Intracellular Fate of *Staphylococcus aureus*

**DOI:** 10.1101/2024.09.13.612871

**Authors:** Marcel Rühling, Fabio Schmelz, Kim Ulbrich, Fabian Schumacher, Julia Wolf, Maximilian Pfefferle, Magdalena Priester, Adriana Moldovan, Nadine Knoch, Andreas Iwanowitsch, Christian Kappe, Kerstin Paprotka, Burkhard Kleuser, Christoph Arenz, Martin J. Fraunholz

**Affiliations:** Chair of Microbiology, Julius-Maximilians-University Würzburg, Theodor-Boveri-Weg, 97074 Würzburg, Germany; Institute of Pharmacy, Freie Universität Berlin, Königin-Luise-Straße 2+4, 14195 Berlin, Germany; Institut for Chemistry, Humboldt Universität zu Berlin, Brook-Taylor-Str. 2, 12489 Berlin, Germany

## Abstract

*Staphylococcus aureus* is an opportunistic pathogen causing severe diseases. Recently, *S. aureus* was recognized as intracellular pathogen, whereby the intracellular niche promotes immune evasion and antibiotic resistance. Interaction of *S. aureus* with versatile host cell receptors was described previously, suggesting that internalization of the pathogen can occur via several pathways. It remains elusive whether the pathway of internalization can affect the intracellular fate of the bacteria. Here, we identified a mechanism governing cellular uptake of *S. aureus* which relies on lysosomal Ca^2+^, lysosomal exocytosis and occurs concurrently to other well-known entry pathways within the same host cell population. This internalization pathway is rapid and active within only few minutes after bacterial contact with host cells. Compared to slow bacterial internalization, the rapid pathway demonstrates altered phagosomal maturation as well as translocation of the pathogen to the host cytosol and ultimately results in different rates of intracellular bacterial replication and host cell death. We show that these alternative infection outcomes are caused by the mode of bacterial uptake.

## Introduction

Sphingolipids are important components of eukaryotic membranes and also serve as bioactive signaling molecules (1). The most abundant sphingolipid, sphingomyelin (SM), resides in the extracellular leaflet of the plasma membrane and hence is exposed to the environment (2, 3). SM is a substrate for sphingomyelinases (SMases) that cleave off the phosphocholine head group to produce ceramide. Acid sphingomyelinase (ASM) is a lysosomal enzyme involved in recycling of SM (4, 5). Several studies demonstrated that ASM can be released by human cells either by Ca^2+^-dependent liberation of lysosomal content via so called lysosomal exocytosis (6–8) or by channeling the enzyme through the Golgi secretory pathway (9, 10). Extracellular ASM is active on the plasma membrane, where it has important roles during repair of membrane wounds (6, 11) but has also been shown to act on the SM-containing low-density lipoprotein (12). The release of ASM through lysosomal exocytosis mediates host cell entry of several human pathogens (7, 8, 13–17).

*Staphylococcus aureus* is a Gram-positive human commensal that asymptomatically colonizes about one third of the human population (18). It is an opportunistic pathogen causing diseases ranging from soft tissue and skin infections (19) to lethal diseases (20, 21). *S. aureus* is notorious for acquiring antibiotic resistances thereby causing >100,000 annual deaths (22). The pathogen possesses intracellular virulence strategies (23, 24) which contribute to antibiotic resistance (25, 26) and immune evasion (27). Internalization of *S. aureus* by host cells is mediated by several adhesins on the staphylococcal surface that bind a plethora of receptors on the host cell plasma membrane (13, 28–38). For instance, staphylococcal fibronectin-binding proteins that form fibronectin bridges to α_5_β_1_ integrins on host cells to initiate internalization (28) or clumping factor B which can interact with the host receptor annexin A2 (31).

After internalization, *S. aureus* resides within a phagosome-like compartment that matures by acquiring proteins associated with early and then late endosomes as well as lysosomes (39–41). In epithelial and endothelial cells, the bacteria translocate to the host cytosol [“phagosomal escape”, (39, 42, 43)]. Phagosomal escape is mediated by so-called phenol-soluble modulins (42), which comprise a family of helical amphiphilic peptides and are transcriptionally controlled by the accessory gene regulator *agr*, a staphylococcal quorum sensing system (44). *S. aureus* replicates in the host cytosol and causes host cell death for example by expression of a cysteine protease. (45).

The mechanism of pathogen entry into host cells has been suggested to dictate the outcome of infections (46–49). For instance, phagosomes formed during natural internalization of *Brucella abortus* by macrophages exhibited different characteristics compared to phagosomes that were artificially generated by phorbol myristate acetate (49). The protozoan parasite *Toxoplasma gondii* actively invades its host cells. Phagosomes generated during this active invasion differed from those that were generated during Fc receptor-mediated uptake of antibody-coated parasites (48). Since previous observation connecting host cell entry and intracellular fate of pathogens were based on artificial induction of internalization, it is unclear whether pathogens can enter host cells within the same population via different naturally occurring mechanisms and if the entry route has a direct influence on the outcome of an intracellular infection.

Here, we describe the internalization of *S. aureus* by tissue cells via a rapid pathway taking place within minutes after contact between bacteria and host cell surface. This pathway requires nicotinic acid adenosine dinucleotide phosphate (NAADP)-dependent mobilization of lysosomal Ca^2+^, followed by lysosomal exocytosis and thereby release of ASM. Since the rapid uptake is concurrent to previously described internalization pathways, it probably has been missed due to long infection times used in traditional infection protocols [e.g., (28, 31, 35, 36)].

*S. aureus* bacteria, which enter host cells of the same cell population via the rapid pathway, cause a distinct infection outcome with altered phagosome maturation, bacterial translocation to host cytosol and eventually host cell death when compared to bacteria that enter host cells at later time points during infection. Thus, the outcome of an infection is decided at the single cell level during bacterial uptake at the host plasma membrane. Bacteria-containing phagosomes that were formed by the ASM-dependent rapid uptake pathway are delayed in their maturation and the bacteria do not escape these organelles efficiently, hence demonstrating a direct link between concurrently active modes of bacterial uptake and the resulting outcomes of intracellular *S. aureus* infection.

## Results

### *S. aureus* triggers lysosomal Ca^2+^ mobilization for host cell invasion

We previously showed that internalization of *S. aureus* by host cells is associated with cellular Ca^2+^ signaling (50). To validate these findings, we treated human microvascular endothelial cells (HuLEC, **Figure 1, A**) or HeLa (**Supp. Figure 1, A**) with the cell permeant Ca^2+^ chelator BAPTA-AM (51) to interfere with host cell Ca^2+^ signaling and subsequently infected the cells with *S. aureus*. We measured the number of internalized *S. aureus* and determined the invasion efficiency by normalizing bacteria numbers detected in treated samples to untreated controls (set to 100%). Invasion efficiency was reduced by BAPTA-AM in a concentration-dependent manner, indicating an involvement of Ca^2+^ in the host cell entry process of *S. aureus*.

**Figure 1.**
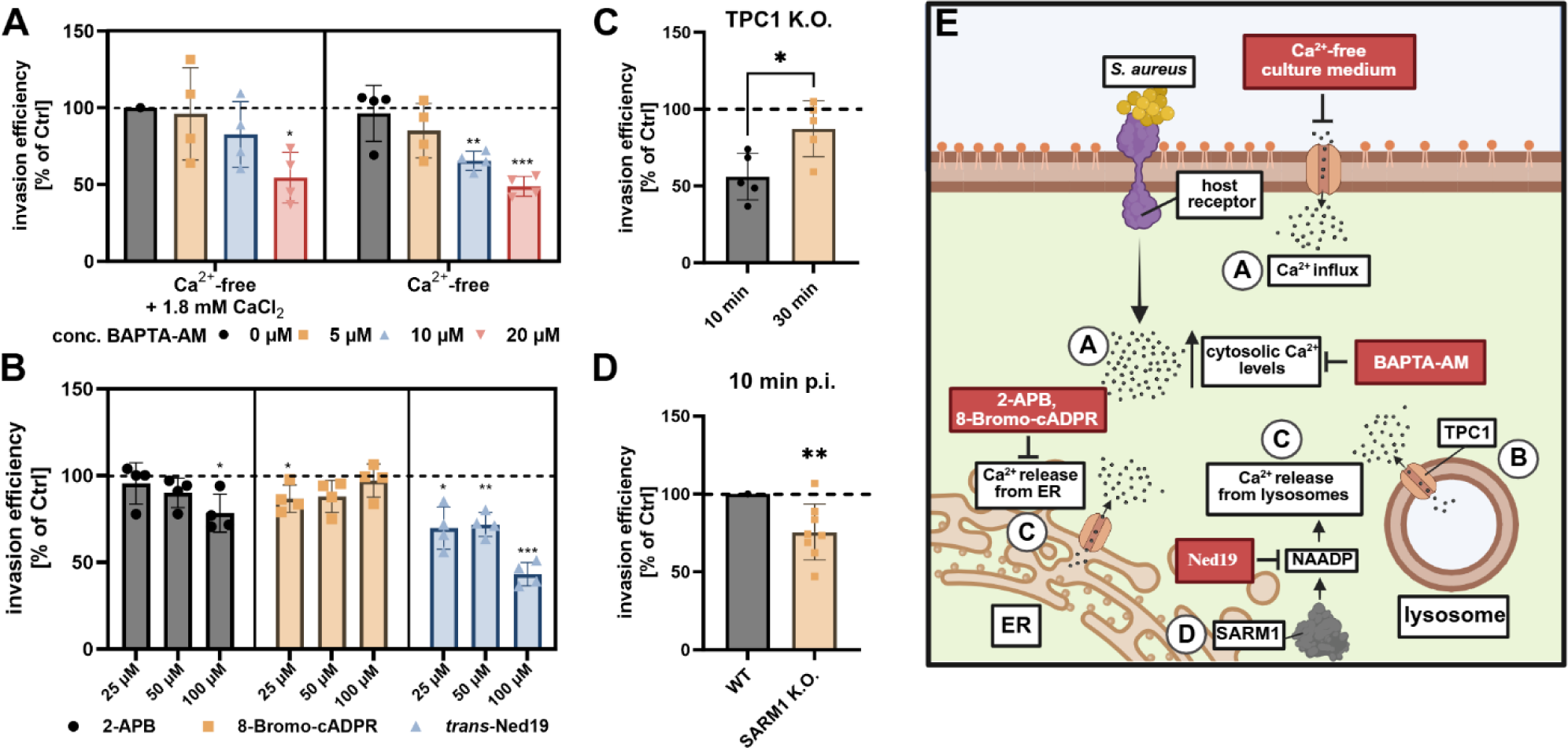
*S. aureus* invasion is dependent on Ca^2+^ liberation from lysosomal stores. Treatment with, BAPTA-AM (A, n≥4), *trans*-Ned19 (B, n≥4), but not 2-APB and 8-Bromo-cADPR (B, n≥4) reduce *S. aureus* internalization by host cells. Genetic ablation of TPC1 (C, n=5) and SARM1 (D, n=8) reduced invasion of *S. aureus* 10 min p.i. HuLEC (A, B), HeLa WT (C, D), HeLa TPC1 K.O. (C) or HeLa SARM1 K.O. (D) cells were treated with the respective substance and were subsequently infected with *S. aureus* JE2 for 30 min if not indicated otherwise. Extracellular bacteria were removed by lysostaphin, and the number of intracellular bacteria was determined by CFU counting. Results were normalized to untreated controls to obtain invasion efficiency (in percent of control). (F) Scheme of host cell Ca^2+^ signaling and interfering agents. Letters indicate figure panels that supported the respective conclusion. Statistics: one sample t-test (A, B,D,), unpaired Student’s t-test (C). Bars represent means ± SD. *p≤0.05, **p≤0.01, ***p≤0.001, ****p≤0.0001.

The cytosolic Ca^2+^ levels, which are crucial for signaling processes, can be elevated by Ca^2+^ influx from the extracellular space. To test whether *S. aureus* internalization is mediated by Ca^2+^ influx, we measured invasion efficiency, in absence of Ca^2+^ from the cell culture medium (**Figure 1, A** and **Supp. Figure 1, A**). Internalization of *S. aureus* by host cells was not affected by absence of Ca^2+^ (“Ca^2+^-free, 0 µM BAPTA-AM”), thereby excluding a Ca^2+^ influx as crucial for *S. aureus* host cell entry. The pore-forming staphylococcal α-toxin was shown to mediate Ca^2+^ influx into host cells (11). *S. aureus* invasion into host cells was largely independent of the production of α-toxin (**Supp. Figure 1, B**), which further supported that the internalization is independent from extracellular Ca^2+^.

Since intracellular Ca^2+^ stores, such as the ER or lysosomes, can serve as alternative source of Ca^2+^ (52), we next blocked Ca^2+^ liberation either from the ER using 2-APB (53) or 8-Bromo-cyclic ADP ribose (8-Bromo-cADPR; (54)) or from lysosomes using *trans*-Ned19 (55). Interference with lysosomal but not ER Ca^2+^ release reduced internalization of *S. aureus* in HuLEC (**Figure 1, B**) or HeLa (**Supp. Figure 1, C, D**) at concentrations which do not affect bacterial viability (**Supp. Figure 1, E** and F).

Ned19 antagonizes NAADP, a second messenger that mediates opening of two pore channels (TPCs), which transport Ca^2+^ from the endo-lysosomal compartment into the cytosol (55). Hence, a HeLa cell pool depleted of TPC1 (**Supp. Figure 1, G**) showed substantially reduced invasion when compared to wildtype HeLa cells. This was particularly pronounced after a 10 min infection pulse, suggesting that Ca^2+^-dependent invasion is important early in infection (**Figure 1, C**).

CD38 is a well-known producer of NAADP in immune cells (56–58). Thus, we tested *S. aureus* invasion in HeLa cells treated with the specific CD38 inhibitor 78c, even though expression of CD38 in these cells is predicted to be very low (proteinatlas.org). We observed no effect on *S. aureus* invasion by CD38 inhibition (**Supp. Figure 1, H**). An alternative NAADP producer, with higher expression in our infection model, is *Sterile Alpha and TIR Motif Containing 1* [SARM1, (59)]. We measured a slightly decreased invasion efficiency 10 min p.i. in a cell pool lacking SARM1 when compared to wildtype cells (**Figure 1, D**; **Supp. Figure 1, I**), suggesting that SARM1 might be involved in NAADP production during internalization of *S. aureus* by host cells. However, there might be other enzymes that produce NAADP (60) during *S. aureus* infection.

Altogether, our findings show that interference with lysosomal Ca^2+^ mobilization limits *S. aureus* invasion particularly early in infection.

### *S. aureus* invasion requires lysosomal exocytosis, ASM and its substrate sphingomyelin on the host cell surface

Since our data suggest that lysosomal Ca^2+^ release facilitates the internalization of *S. aureus* by host cells, we speculated that lysosomal exocytosis is involved in bacterial invasion. Lysosomal exocytosis is a Ca^2+^-dependent process during which lysosomes fuse with the plasma membranes and release their content. The internalization of several human pathogens has been shown to depend on lysosomal exocytosis (8, 16, 61–63).

To visualize lysosomal exocytosis, we developed an assay that makes use of the split NanoLuc luciferase system [**Figure 2, A**; (64, 65)]. We here engineered HeLa cells to express the high-affinity NanoLuc peptide HiBiT between the signal peptide and the mature chain of lysosomal-associated membrane protein 1 (LAMP1). Transient localization of LAMP1 to the cell surface of mammalian cells is a hall mark for lysosomal exocytosis (66). To measure the release of the HiBiT-tagged LAMP1, we added LgBit together with the NanoLuc substrate Furimazine to the cell culture medium and measured luminescence of reconstituted NanoLuc in a microplate reader. Maximum luminescence was detected ∼10-15 min upon starting the assay and decline afterwards, likely due to exhaustion of the luciferase substrate (**Figure 2, B**). Treatment with the Ca^2+^ ionophore ionomycin, a known activator of lysosomal exocytosis (11, 67), increased the luminescence, whereas Vacuolin-1, an inhibitor of lysosomal exocytosis (68), decreased luminescence when compared to an untreated control. Thus, we concluded that our assay successfully quantified exposure of LAMP1 on the host cell surface.

**Figure 2.**
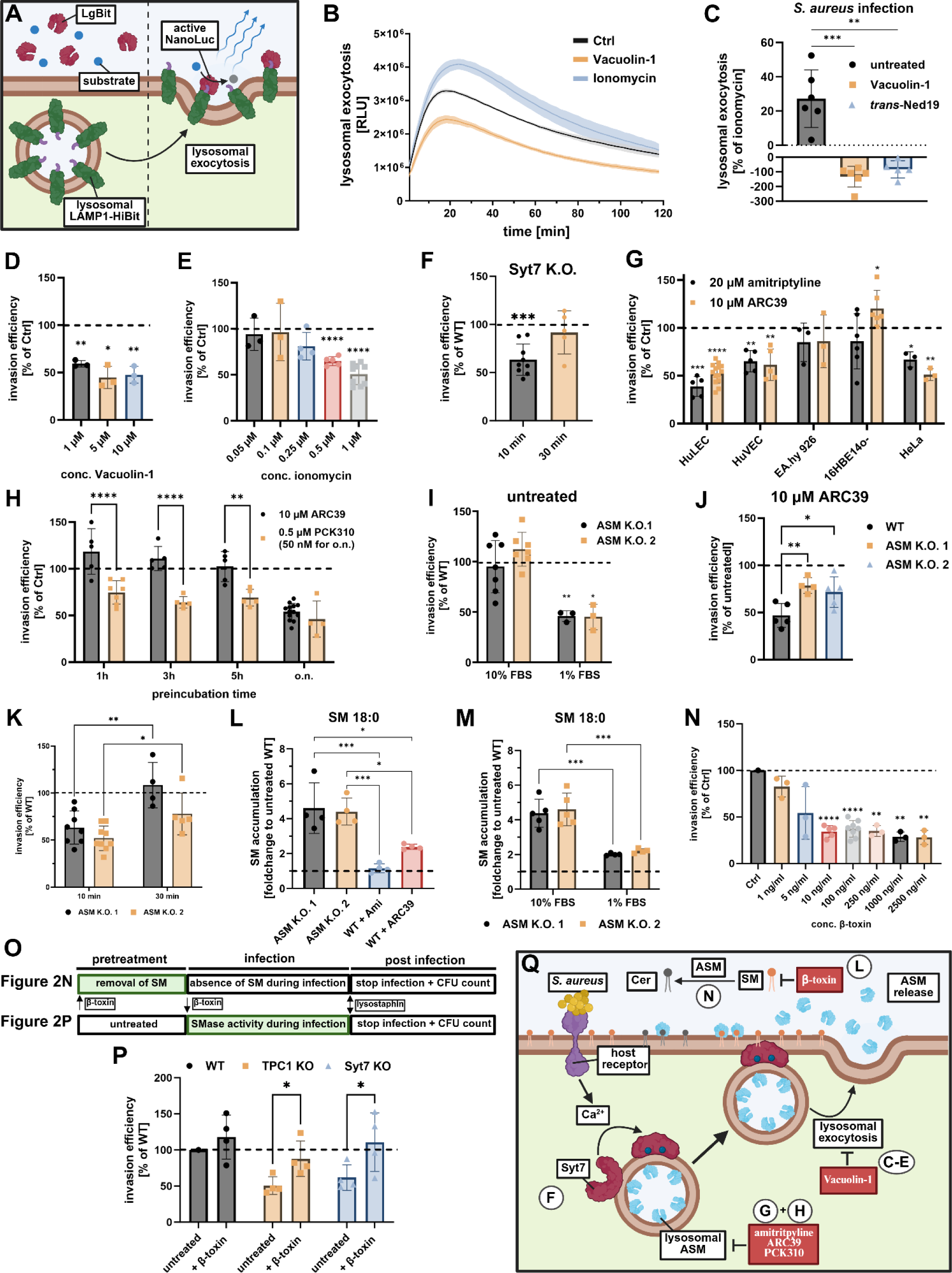
*S. aureus* invasion requires lysosomal exocytosis, ASM and plasma membrane sphingomyelin. **(A, B) Luminescence-based lysosomal exocytosis assay.** HeLa cells expressing LAMP1-HiBit were treated with 1 µM ionomycin (immediately before the measurement) or 1 µM Vacuolin-1 (75 min before the measurement) in presence of LgBit and NanoLuc substrate. Lysosomal exocytosis was monitored by measuring chemiluminescence in a Tecan microplate reader. n=3 **(C) *S. aureus* triggers lysosomal exocytosis during infection.** HeLa cells expressing LAMP1-HiBit were pretreated for 75 min with 1 µM Vacuolin-1 or 200 µM *trans*-Ned19. Subsequently, cells were infected with *S. aureus* JE2 (MOI10), treated with 1 µM ionomycin or left untreated and lysosomal exocytosis was determined by measuring chemiluminescence in a Tecan microplate reader 10 min p.i. Luminescence detected in infected samples was scaled to untreated controls (0 %) and ionomycin (100%) to determine the proportion of all “releasable” lysosomes liberated during infection. n=6. **(D-F) *S. aureus* invasion requires Syt7-dependent lysosomal exocytosis.** HuLEC were treated with the lysosomal exocytosis inhibitor Vacuolin-1 or the lysosomal exocytosis inducer ionomycin and invasion efficiency of *S. aureus* was determined 30 min p.i. (D, E, n≥3). Invasion efficiency of *S. aureus* JE2 was determined in HeLa Syt7 K.O. cells 10 min and 30 min p.i. (F, n≥5; data normalized to HeLa wildtype). **(G, H) *S. aureus* internalization is dependent on ASM activity in endothelial and epithelial cells.** Indicated cell lines were treated with the ASM inhibitors amitriptyline or ARC39 (G) and invasion efficiency of *S. aureus* was determined (n≥3). HuLEC were treated with PCK310 or ARC39 for the indicated periods. Subsequently, cells were infected with *S. aureus* for 30 min and invasion efficiency was determined by CFU counting (H). n≥4. **(I) *S. aureus* internalization by ASM-deficient cells is only affected when cells were cultured in medium containing low FBS concentrations.** ASM-deficient and wildtype HeLa cells were either cultured in 1% or 10% FBS prior to infection. Number of internalized bacteria was measured 10 min p.i.. Invasion efficiency was determined by normalizing the results of ASM-deficient cells to wildtype cells cultured in the corresponding FBS concentration. n≥3. **(J) Genetic ablation of ASM renders *S. aureus* invasion insensitive towards ARC39.** HeLa ASM K.O. cells were cultured in 10% FBS and treated with 10 µM ARC39 or were left untreated and the number of invaded bacteria was determined 30 min p.i.. Invasion efficiency was determined by normalization to corresponding untreated samples. n≥4 **(I) Absence of ASM has a stronger effect on *S. aureus* internalization early during infection.** ASM-deficient and wildtype HeLa cells were cultured in 1% FBS prior to infection. Invasion efficiency was determined 10 min and 30 min p.i. n≥4. **(L) Genetic ablation of ASM is accompanied by massive alteration in cellular sphingolipid profiles.** HeLa wildtype and ASM K.O. cells were treated with 20 µM amitriptyline (75 min) and 10 µM ARC39 (22h) or left untreated, as indicated. Whole cell sphingolipid profiles were measured by HPLC-MS/MS and foldchange of SM accumulation was determined relative to untreated wildtype cells (set to 1). Here exemplified by SM 18:0, for SM species with other acyl chain length see **Supp. Figure 3, C**. n= 4. **(M) SM accumulation is less severe when ASM-deficient cells are cultured in 1% FBS.** HeLa wildtype and ASM K.O. cells were either cultured in 1% or 10% FBS. Whole cell sphingolipid profiles were measured by HPLC-MS/MS and foldchange of SM accumulation in ASM K.O. cells were determined relative to untreated wildtype cells (set to 1). Here exemplified by SM 18:0, for SM species with other acyl chain length see **Supp. Figure 3, D**. (N) *S. aureus* invasion requires SM on host cell plasma membranes. HuLEC were treated with the bacterial SMase β-toxin for 75 min and were subsequently infected with *S. aureus* JE2 for 30 min (n≥3). (O) Experimental design for β-toxin treatment of host cells during *S. aureus* infection. Host cells were either pretreated with β-toxin to remove SM from the plasma membrane prior to infection (upper panel; Figure 2**, N).** Alternatively, β-toxin was added together with the bacteria to rescue the absence of ASM (lower panel, Figure 2**, P)**. **(P) Presence of extracellular SMase activity restores the invasion defect in TPC1 and Syt7 K.O. cell lines.** HeLa wild type as well as TPC1 or Syt7 KO cell lines were infected with *S. aureus* JE2 in presence of 100 ng/ml of the bacterial SMase β-toxin and invasion efficiency 10 min p.i. was determined (n=4). In each experiment the numbers of intracellular *S. aureus* JE2 were determined by lysostaphin protection assay and CFU counting. Results obtained for the tested condition were normalized to the wildtype cell line or the mock-treated control. (**Q) Scheme of lysosomal exocytosis and ASM release as well as interfering agents.** Cer: ceramide, SM: sphingomyelin, ASM: acid sphingomyelinase. Statistics: one sample t-test (D-G, L), one-way ANOVA and Dunnett’s multiple comparisons test (C, J). Mixed-effects model (REML) and Šídák’s multiple comparisons (H). Two-way ANOVA and Tukey’s multiple comparison (K). Two-way RM ANOVA and Šídák’s multiple comparison (N) Bars represent means ± SD. *p≤0.05, **p≤0.01, ***p≤0.001, ****p≤0.0001.

Next, we monitored LAMP1 surface levels during *S. aureus* infection. We assumed that ionomycin would trigger exocytosis of all “releasable” lysosomes. Hence, the detected luminescence signal during infection was scaled to an untreated control (=0 %) as well as ionomycin (=100%), to express LAMP1 surface levels as percentage of all “releasable” lysosomes. We detected that ∼30% of all “releasable” lysosomes were liberated from the reporter cells 10 min after infection with *S. aureus* (**Figure 2, C**). Pretreatment of host cells with Vacuolin-1 or Ned-19 strongly reduced LAMP1 surface levels, suggesting that *S. aureus* triggers lysosomal exocytosis in an NAADP-dependent manner during invasion.

To test whether lysosomal exocytosis is required for *S. aureus* internalization by host cells, we measured invasion efficiency in HuLEC (**Figure 2, D**) and HeLa cells (**Supp. Figure 2, A**) pretreated with Vacuolin-1. Invasion was strongly reduced in inhibitor-treated samples, while the survival of the bacteria was not affected by the compound (**Supp. Figure 2, B**).

Next, we triggered lysosomal exocytosis in HuLEC by addition of ionomycin prior to infection, thereby depleting the pool of “releasable” lysosomes and thus preventing lysosomal exocytosis during infection. Ionomycin pretreatment reduced invasion efficiency in a concentration-dependent manner (**Figure 2, E**). Bacterial growth and survival as well as host cell integrity were not affected at the concentrations used (1 µM, **Supp. Figure 2, C-E**).

Synaptotagmin 7 (Syt7) is known to support fusion of lysosomes with the plasma membrane in a Ca^2+^-dependent manner (69). Accordingly, Syt7-depleted HeLa cells demonstrated lower bacterial invasion (**Figure 2, F**). As already observed for TPC1 (**Figure 1, C**), this was most pronounced early in infection. Taken together, our results support the hypothesis that lysosomal exocytosis is important for invasion of host cells by *S. aureus* particularly early during infection.

Lysosomal exocytosis results in release of ASM (6, 11), an enzyme which has previously been associated with the internalization of several bacterial and viral pathogens (8, 13, 16, 62, 63, 70) and thus, we tested if ASM activity is also required for the uptake of *S. aureus* by host cells. Therefore, we treated different cell lines prior to infection with amitriptyline, a functional inhibitor of ASM (FIASMA), or the competitive ASM inhibitor ARC39 (71) and infected host cells with *S. aureus*. While invasion in HuLEC, human umbilical vein endothelial cells (HuVEC) and HeLa was reduced by both inhibitors, invasion in bronchial epithelial 16HBE14o^-^ and the endothelial-epithelial hybrid cell line EA.hy926 was not or only marginally altered (**Figure 2, G**). The used inhibitor concentrations did not affect bacterial survival (**Supp. Figure 2, F, G**). Furthermore, we excluded that differences among cell lines arose from differential ASM activity or sensitivity to the inhibitors by monitoring enzymatic activity [**Supp. Figure 2**, H; (72) and **Supp. Figure 2, I**; (73)].

Next, we treated HuLEC with 10 µM ARC39 or 0.5 µM PCK310, a fast-acting ASM inhibitor. Already after 1h preincubation, PCK310 reduced invasion of *S. aureus*, whereas ARC39 required overnight treatment for a similar reduction (**Figure 2, H**). We confirmed the reduction of internalized colony forming units (CFU) by amitriptyline and PCK310 treatment with a microscopy-based approach, where we determined the number of invaded fluorescent bacteria per host cell (**Supp. Figure 2, J**).

Next, we generated two ASM-depleted HeLa cell pools (referred to as ASM K.O. 1 and 2) using CRISPR/Cas9 (74) with two distinct small guide RNAs (sgRNAs). We confirmed successful ablation of cellular ASM activity in both ASM-depleted cell lines [(73), **Supp. Figure 2, K**].

We did not detect a reduced invasion in ASM K.O. cells, when we infected in common infection medium containing 10% heat-inactivated fetal bovine serum (FBS; **Figure 2, I**).

To test if unspecific side effects of the ASM inhibitors could have contributed to reduced invasion in our previous experiments, we treated wildtype and ASM K.O.s with ARC39 and tested for *S. aureus* invasion efficiency (**Figure 2, J**). In wildtype cells, bacterial invasion again was strongly reduced after ARC39 treatment, whereas only a slight reduction was observed in the K.O.s, which may represent unspecific side effects of the inhibitor. Since the reducing effect of ARC39 on invasion is almost completely lost in ASM K.O. cells, the reduced invasion in wildtype cells upon ARC39 treatment predominantly must be caused by the inhibition of ASM.

FBS contains high levels of SM (75) as well as ASM (76), even though the enzymatic activity is strongly reduced upon heat inactivation (11).

To test whether FBS confounded our invasion experiments, we cultivated WT as well as ASM K.O. cells in medium with reduced FBS concentration (1%) and determined the *S. aureus* invasion efficiency (**Figure 2, I**). In both ASM-depleted cell lines, we detected reduced *S. aureus* invasion when compared to wildtype cells only when cells were cultured in 1% FBS. Similar to our observations for TPC1 and Syt7 K.O. cells, the absence of ASM had a more pronounced effect on host cell entry of *S. aureus* early during infection, if cells were cultured in 1% FBS (**Figure 2, K**).

ASM activity in FBS could be taken up by ASM K.O. cells, thereby potentially complementing cellular ASM activity and restoration of *S. aureus* invasion. However, the FBS concentration did not affect cellular ASM activity of ASM K.O. cells (**Supp. Figure 2, L**).

Absence of ASM results in cellular accumulation of SM and usually is accompanied by the neurodegenerative diseases, Niemann-Pick syndrome A and B (77). Hence, we compared the sphingolipidome of inhibitor-treated wildtype and ASM K.O. cells that we either cultured in 1% or 10% FBS by high-pressure liquid chromatography coupled to tandem mass spectrometry (HPLC-MS/MS). Cellular SM levels were determined by ratios of SM vs. ceramide concentrations (ASM educt vs. product) for lipid species with different acyl chain lengths (**Supp. Figure 3, A** and B). To assess the SM accumulation upon ASM K.O. or inhibitor treatment, the fold changes of the detected SM levels compared to untreated WT cells were determined (set to 1, **Supp. Figure 3, C** and D). Whereas we did observe a moderate SM accumulation upon ARC39 treatment, e.g. for SM 18:0 (**Figure 2, L**) but also other SM species (**Supp. Figure 3, A** and C), cells exposed to amitriptyline only showed a slight increase in SM 16:0, which can be explained by the shorter preincubation time used for amitriptyline (75 min vs. 22h; **Supp. Figure 3, A** and C). However, if ASM K.O. cells were cultured in 10% FBS, we observed markedly increased SM levels when compared to either untreated or inhibitor-treated wildtype cells [e.g. for SM 18:0 (**Figure 2, L**) and other SM species (**Supp. Figure 3, A** and C)]. The relatively low increase of SM levels after inhibitor treatment likely arises from short inhibitor pulses (in the range of minutes or hours), whereas SM in ASM K.O. cell lines accumulates over the entire culture period (days or weeks since gene ablation). Strikingly, ASM K.O. cells accumulated markedly less SM 18:0 (**Figure 2, M**), SM 16:0 and SM 20:0 (**Supp. Figure 3, D**), when cultured in 1% FBS, suggesting that FBS is an important source of SM in absence of ASM.

**Figure 3.**
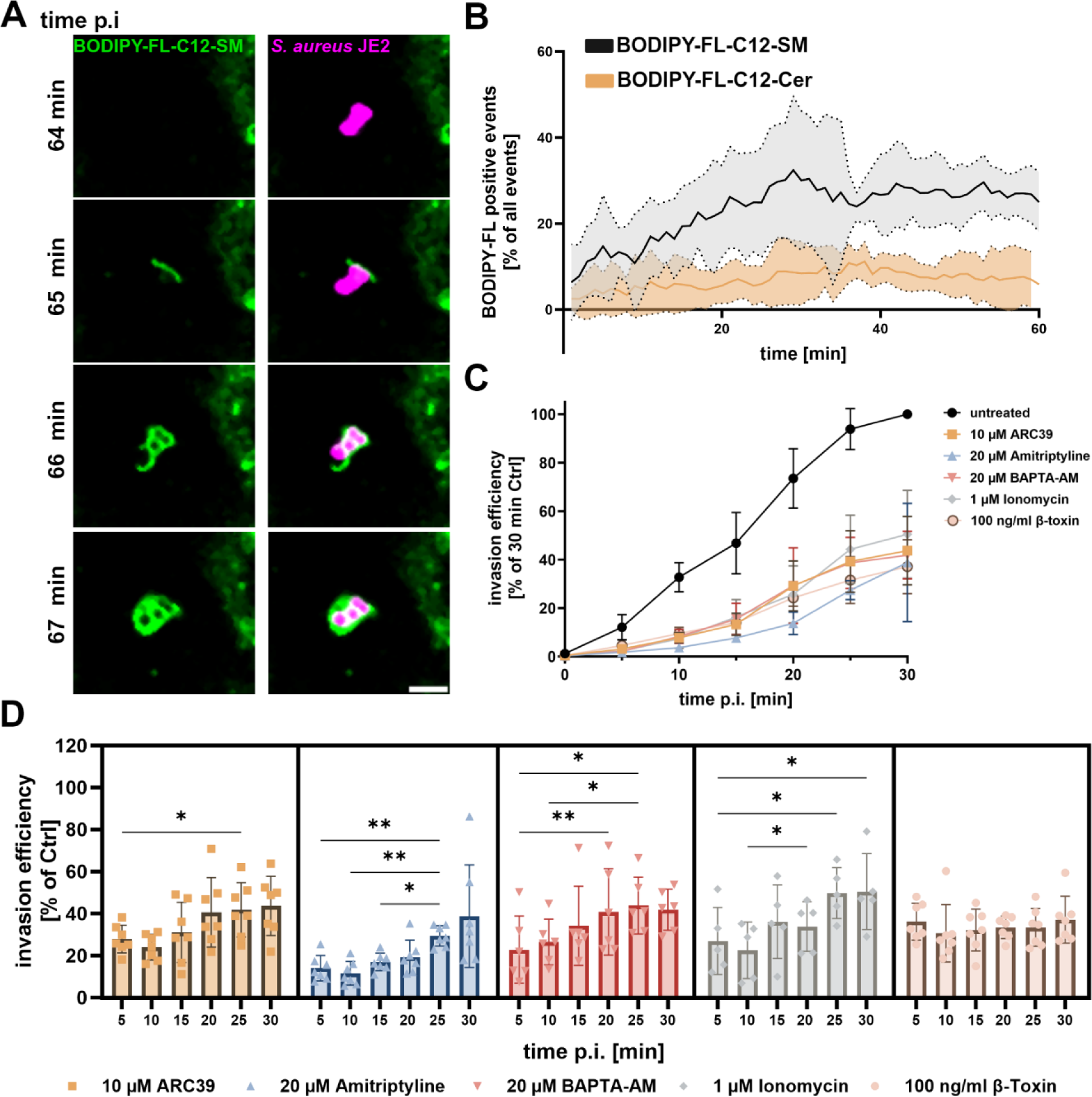
ASM- and Ca^2+^-dependent uptake is rapid and predominantly mediates invasion early in infection. **(A, B) *S. aureus* associates with SM early during invasion.** HuLEC were pretreated with BODIPY-FL-C12-SM or BODIPY-FL-C12-ceramide and infected with red-fluorescent *S. aureus.* Scale bar: 5 µm. Infection was monitored by live cell imaging and the proportion of bacteria that associated with the lipid analogs was quantified in each individual frame (SM: n=5, Cer: n=3). (**C, D) ASM and Ca^2+^-dependent invasion is rapid.** HuLEC were treated with ARC39, amitriptyline, BAPTA-AM, ionomycin, or the bacterial SMase β-toxin. Then, cells were infected with *S. aureus* and the number of invaded bacteria was determined after different time periods. The results were either normalized to the 30 min time point of untreated controls (C) or to the corresponding time points of the untreated controls (D). (n ≥5). Statistics: Mixed effect analysis and Tukey‘s multiple comparison. Graphs represent means ± SD. *p≤0.05, **p≤0.01, ***p≤0.001, ****p≤0.0001.

In summary, we detected moderate SM accumulation in conditions (ASM inhibitor treatment or ASM K.O.s cultured in 1% FBS) where we also detected reduced *S. aureus* invasion and a strong SM accumulation (ASM K.O. cultured in 10% FBS) when the number of internalized bacteria was not affected when compared to untreated wildtype cells.

We hypothesize that strong cellular SM accumulation generally increases ASM-independent host cell invasion of *S. aureus*, while the ASM-dependent uptake pathway is absent. This is supported by the high residual internalization of *S. aureus* by ASM K.O. cells cultured in 10% FBS upon ARC39 treatment. Previously, it was shown that *S. aureus* host cell entry is limited by caveolin-1, a protein associated with lipid microdomains (78, 79). Absence of ASM was demonstrated to interfere with caveolin-1-associateded endocytosis (80, 81) and thus, the increased invasion in ASM K.O. cells may be caused by dysfunctional caveolin-1.

Taken together, our results show that ASM is involved in *S. aureus* invasion of host cells in a cell-type specific manner.

Release of ASM by host cells results in cleavage of SM on the host cell surface, a process that previously was suggested to facilitate cell entry of human pathogens (7, 8, 13–17). To test whether *S. aureus* invasion requires SMase activity on the host cell surface, we removed SM, the substrate of ASM, from plasma membranes of HuLEC (**Figure 2, N**) and HeLa (**Supp. Figure 2, M**) by pretreatment with the bacterial SMase β-toxin (see **Figure 2, O**). We confirmed β-toxin-dependent SM conversion in living cells by detecting the ceramide metabolites of fluorescent SM analogs [**Supp. Figure 2**, N; (82)]. Plasma membrane pretreatment with bacterial SMase reduced *S. aureus* invasion efficiency in a concentration-dependent manner in both cell lines. 10 ng/ml β-toxin were sufficient to decrease invasion to ∼30% of that measured in untreated control cells. Interestingly, a 250-fold higher concentration did not lead to further reduction, suggesting that 10 ng/ml β-toxin were sufficient to quantitatively convert SM to ceramide at the cell surface. In line with our previous experiments, this suggests that multiple *S. aureus* invasion pathways exist within the same host cell population, of which at least one requires SM and its enzymatic breakdown on the plasma membrane.

Since we detected enhanced Ned19-sensitive lysosomal exocytosis during *S. aureus* infection (**Figure 2, C**), we hypothesized that absence of TPC1 or Syt7 should impair lysosomal exocytosis and thereby release of ASM. Therefore, we infected wildtype as well as TPC1 or Syt7 K.O. cells with *S. aureus* in presence (but without pretreatment, see **Figure 2, O**) of 100 ng/ml β-toxin and determined the number of intracellular bacteria after 10 min (**Figure 2, P**). The addition of β-toxin to the culture medium completely rescued the invasion defect of the K.O. cells. This indicates that reduced bacterial internalization in absence of TPC1 and Syt7 results from a lack in ASM delivery to the cell surface by Ca^2+^-dependent lysosomal exocytosis and that absence of ASM from the host cell surface, can be compensated by exogenous addition of the bacterial SMase β-toxin.

We want to emphasize the difference in experimental application of β-toxin (**Figure 2, O**) where we i) pretreat host cells with the bSMase to remove SM from the plasma membrane (**Figure 2, N** and **Supp. Figure 2, M**) and ii) rescue the absence of exocytosed ASM from the host cell surface by exogenous addition of β-toxin during the infection (**Figure 2, P**). In the former case, pretreatment leads to removal of SM from the plasma membrane thus preventing SM conversion by ASM during infection and resulting in reduced invasion. The latter leads to SM conversion by the bacterial SMase during infection thereby increasing invasion.

In summary, our data suggest the existence of a *S. aureus* invasion pathway that depends on the metabolic breakdown of SM by ASM on the host cell surface (**Figure 2, Q**).

### ASM- and Ca^2+^-mediated invasion is rapid

To study the role of SM in *S. aureus* uptake kinetics, we incubated HuLEC with fluorescent sphingolipid analogs and recorded *S. aureus* infection by time lapse imaging [**Figure 3, A**; **Supp. Video 1**; **Supp. Figure 4**; **Supp. Video 2**; (73)]. We observed that the bacteria were rapidly engulfed by SM-containing membrane compartments. While association of bacteria with BODIPY-FL-C12-SM increased over time during the first 30 min of infection, this was not observed for BODIPY-FL-C12-Ceramide, suggesting that *S. aureus* invasion specifically relies on interaction with SM in host cell plasma membranes (**Figure 3, B**).

**Figure 4.**
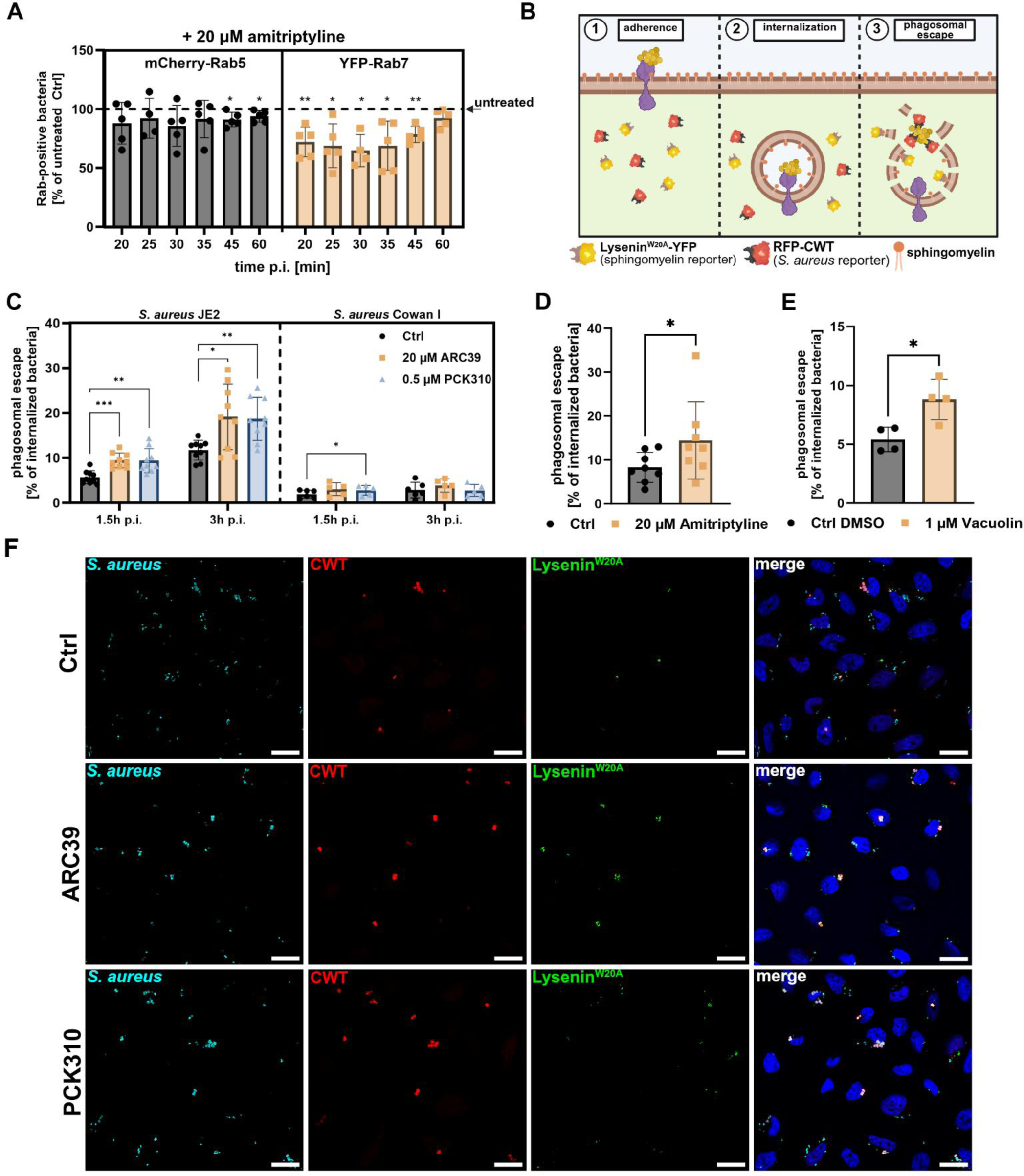
Blocking ASM-dependent invasion affects phagosomal maturation and escape during *S. aureus* infection. **(A) Inhibition of ASM delays formation of Rab7-positive phagosomes.** HeLa cells expressing mCherry-Rab5 and YFP-Rab7 were either treated with amitriptyline or left untreated. Then, cells were infected with *S. aureus* JE2 for indicated periods. Extracellular bacteria were removed and percentage of intracellular bacteria which associated with Rab5 or Rab7 was determined by CLSM. Results obtained for amitriptyline-treated samples were normalized to untreated controls (n=5). **(B) Detection of phagosomal SM and phagosomal escape by a reporter cell line expressing Lysenin^W20A^-YFP and RFP-CWT.** After internalization, *S. aureus* resides in a phagosome preventing the recruitment of RFP-CWT to *S. aureus* cell wall and Lysenin^W20A^-YFP to luminal SM, respectively. When the bacteria lyse the phagosomal membrane, luminal SM gets exposed to the cytosol and attracts Lysenin^W20A^-YFP, while RFP-CWT is recruited to the staphylococcal surface. (**C-F) Blocking ASM-dependent internalization affects phagosomal escape.** RFP-CWT and Lysenin^W20A^-YFP expressing HeLa were treated with PCK310,ARC39 (C, F), amitriptyline (D) or Vacuolin-1 (E) and infected with *S. aureus* strains JE2 or Cowan I. By CLSM, proportions of bacteria that recruited RFP-CWT (phagosomal escape) were determined 3h p.i. if not indicated otherwise (n≥3). **Statistics:** one sample t-test (A), Mixed effects analysis (REML) and Tukey’s multiple comparison (C,), paired Student’s t-test (D, E) Graphs represent means ± SD. *p≤0.05, **p≤0.01, ***p≤0.001, ****p≤0.0001.

Next, we determined the invasion efficiency of *S. aureus* in a time-dependent manner and blocked invasion by either ASM inhibitors, ionomycin, BAPTA-AM or β-toxin pretreatment. We found a time-dependent increase in intracellular bacteria, whereby all treatment conditions reduced bacterial invasion at every time point with the untreated control at 30 min set to 100% (**Figure 3, C**). We next normalized the data to the untreated control of the corresponding time point. Except for the β-toxin treatment, the effect of all other treatments was most pronounced early (5-10 min) after infection (**Figure 3, D**). For instance, ionomycin treatment massively reduced *S. aureus* invasion to ∼20% of untreated controls when we infected for 10 min. We concluded that in untreated controls 80% of the bacteria entered host cells via the pathway that depends on lysosomal exocytosis, Ca^2+^ and ASM, which is blocked by ionomycin treatment. Consequently, bacteria that were able to invade host cells in ionomycin-treated samples (the residual ∼20%) must have employed other pathways. By contrast, in samples infected for 30 min a higher proportion of bacteria was able to enter host cells despite ionomycin treatment (∼50% compared to untreated controls). This suggests that only 50% of all invaded bacteria were internalized in an ASM-and Ca^2+^-dependent fashion upon the longer infection pulse. Together with the observed invasion defects in host cells with gene deletions in TPC1 (**Figure 1, C**), Syt7 (**Figure 2, F**) and ASM (**Figure 2, K**), which we exclusively detected during short infection times, our data suggest that early during infection bacteria enter host cells predominantly in an ASM- and Ca^2+^-dependent manner. Thus, we concluded that the internalization pathway that depends on lysosomal exocytosis, ASM and Ca^2+^ is rapid when compared to other host cell entry pathways.

### The mode of *S. aureus* invasion affects infection outcome

Next, we tested whether the pathway used for host cell entry also affects the intracellular fate of *S. aureus*. After invasion, the bacteria reside within Rab5-positive early and subsequently, in Rab7-positive late phagoendosomes (39). To trace phagosome maturation, we generated a HeLa cell line stably expressing mCherry-Rab5 as well as YFP-Rab7 and infected with *S. aureus*. We observed that the bacteria transiently associate with Rab5 for a few minutes, before they become positive for Rab7 (**Supp. Video 3**).

We next blocked the ASM-dependent invasion pathway by amitriptyline treatment and infected with *S. aureus* for different time periods. We determined the proportion of bacteria that was associated with Rab5 and/or Rab7 (**Figure 4, A**; see **Supp. Figure 5** for bacteria proportion that associates with Rab7). While ASM inhibition only marginally affected *S. aureus* association with Rab5-positive phagosomes, the proportion of bacteria residing in Rab7-positive vesicles was significantly reduced when compared to the untreated control. This observation was restricted to shorter infection pulses (5-45 min) and was not observed for long infection periods (60 min).

**Figure 5.**
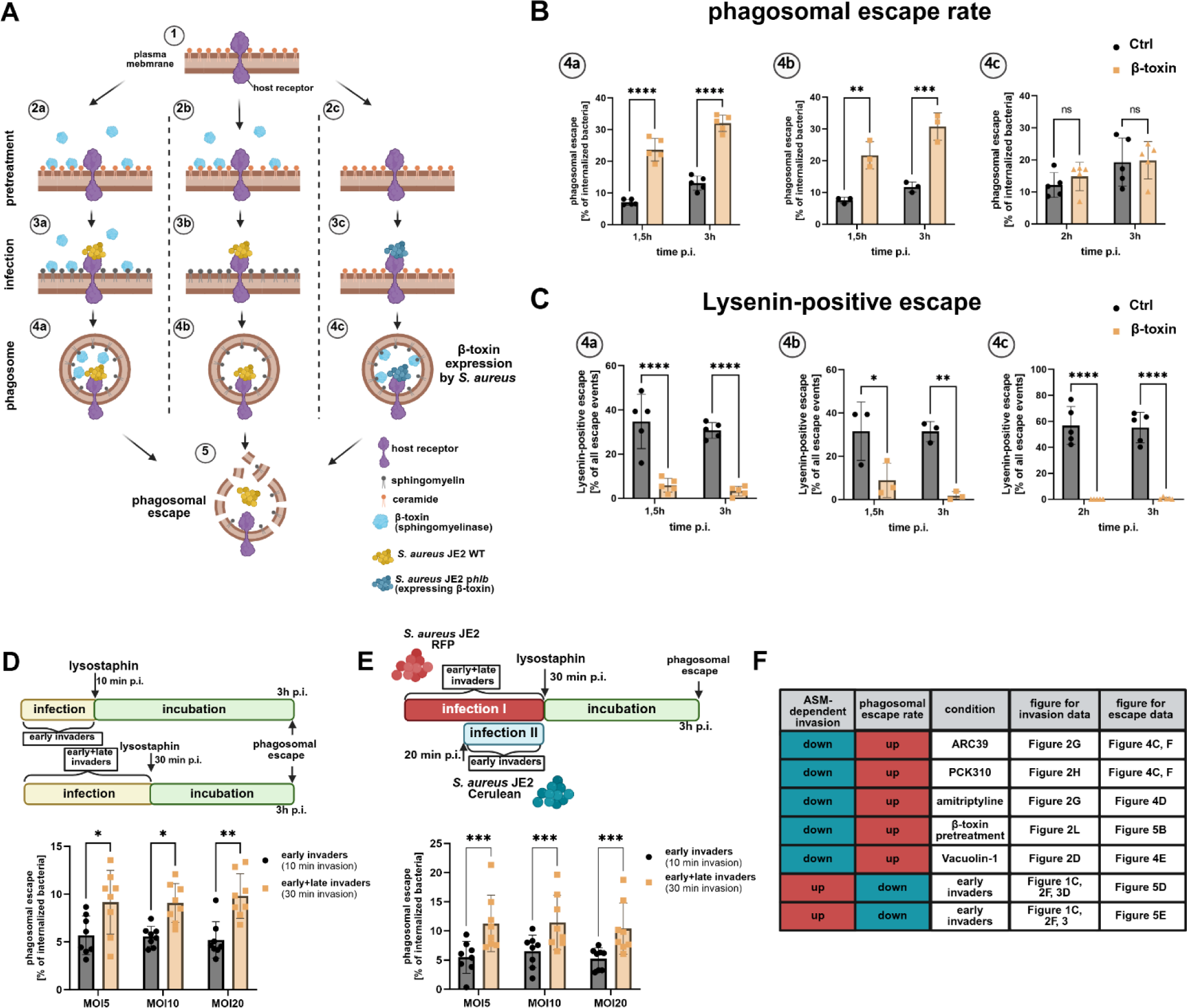
The intracellular fate of *S. aureus* is determined by host cell entry. **(A-C) Phagosomal escape depends on presence of plasma membrane SM during invasion, but not presence of SM within phagosomal membranes** HeLa RFP-CWT Lysenin^W20A^-YFP were pretreated with β-toxin to remove surface SM (2a, 2b) or left untreated (2c). Then, cells were infected with *S. aureus* JE2 in presence (3a) or absence (3b) of β-toxin. Untreated samples (3c) were infected with *S. aureus* JE2 harboring a plasmid either encoding β-toxin and the fluorescence protein Cerulean (pCer+*hlb*) or solely Cerulean (pCer). Proportion of bacteria that recruited RFP-CWT (phagosomal escape, B) and the percentage of phagosomal escape events that additionally were positive for Lysenin^W20A^-YFP (C) were determined at indicated time points p.i. (n=5). **(D, E) Early ASM-dependent invaders possess lower escape rates than late invaders.** HeLa cells expressing RFP-CWT were infected with indicated MOIs of *S. aureus* JE2 either for 10 min (early invaders) or 30 min (early+late invaders). Phagosomal escape rates were determined 3h p.i. (D). HeLa reporter cells expressing YFP-CWT were infected with an MOI=5 of *S. aureus* JE2 expressing a fluorescent protein (e.g. RFP) for 30 min (early+late invaders). After 20 min, the same samples were infected with *S. aureus* JE2 expressing another fluorophore (e.g. Cerulean) for 10 min (early invaders) and phagosomal escape was determined 3 h p.i. (E). (F) Summary of experiments analyzing the influence of invasion pathway on phagosomal escape. The measured effects of different conditions on rapid ASM-dependent invasion and accompanied alterations in phagosomal escape rates are summarized. red = increased, blue=decreased. **Statistics:** Two-way ANOVA and Šídák’s multiple comparisons test (B-E). Graphs represent means ± SD. *p≤0.05, **p≤0.01, ***p≤0.001, ****p≤0.0001.

We also observed a reduced proportion of bacteria associated with Rab7-positive membranes after treatment with PCK310 or bacterial SMase (**Supp. Figure 5, B, C**). Forty-five minutes p.i., about half of the bacteria were localized to Rab7-positive vesicles in untreated cells, whereas only ∼40% associated with Rab7 upon amitriptyline, PCK310 or β-toxin treatment (**Supp. Figure 5, C**). However, this was not caused by translocation of *S. aureus* to the host cytosol, which was excluded by expressing the fluorescence reporter RFP-CWT (42) in the YFP-Rab7 cell line. Accordingly, RFP-CWT-labelled bacteria in the host cytosol were excluded from the data set [**Supp. Figure 5**, B-D,, (42)]. Hence, ASM-dependently generated phagoendosomes possess different maturation dynamics when compared to phagosomes formed in an ASM-independent fashion.

Next, we measured phagosomal escape while simultaneously detecting SM content of disrupted phagosomal membranes. Therefore, we used a HeLa reporter cell line that cytosolically expresses the phagosomal escape reporter RFP-CWT (42) as well as the SM reporter Lysenin^W20A^-YFP (83), respectively (**Figure 4, B**). To validate the reporter system, we removed SM from the plasma membrane by β-toxin treatment prior to infection and monitored phagosomal escape by live cell imaging (**Supp. Figure 6, Supp. Video *4***). Bacterial translocation into the host cytosol is, again, indicated by recruitment of RFP-CWT. In untreated cells, phagosomal escape was accompanied by recruitment of Lysenin^W20A^-YFP to vesicular membranes, suggesting that SM located at the luminal leaflet of phagosomes was exposed to the cytosol. By contrast, Lysenin^W20A^-YFP was not recruited to escape events in β-toxin-treated cells, indicating that the SMase had removed SM from the membranes.

**Figure 6.**
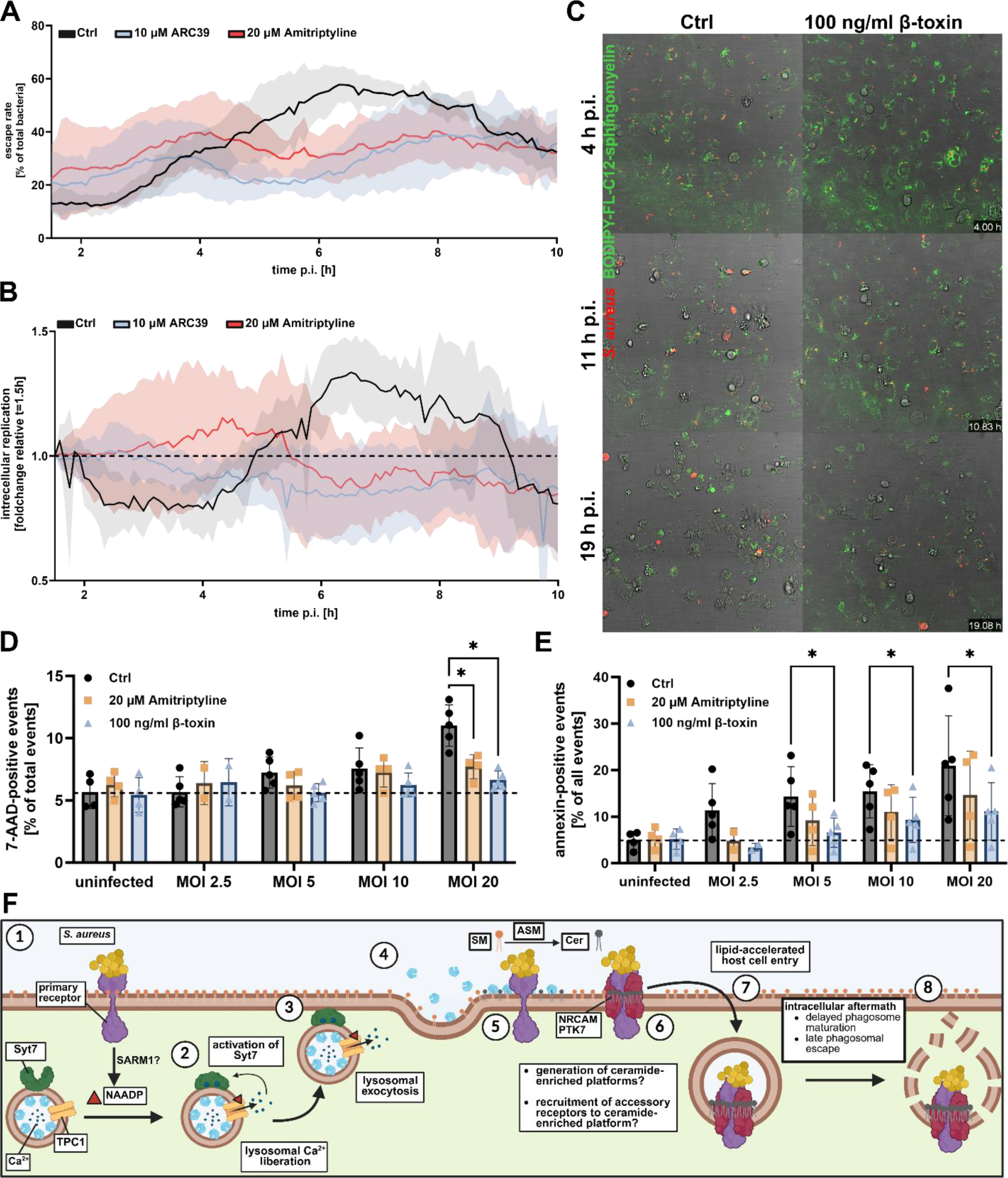
Blocking ASM-dependent internalization affects intracellular replication and host cell survival. **(A,B)** HeLa cells expressing RFP-CWT and Lysenin^W20A^-YFP were treated with amitriptyline or ARC39 or were left untreated (Ctrl). The cells were subsequently infected with *S. aureus* and infection was monitored for 10 h by CLSM. Proportion of escaped bacteria (A) or the intracellular replication (B) was determined (n= 5). **(C)** HuLEC were stained with BODIPY-FL-C12-SM (green), and were either pretreated with 100 ng/ml of β-toxin or were left untreated. Then, cells were infected with *S. aureus* (red) and infection was monitored by CLSM for 19 h. **(D, E)** HuLEC were treated with amitriptyline or β-toxin and subsequently infected with *S. aureus*. After 21 h, plasma membrane integrity was measured by 7-AAD staining (D), and proportion of apoptotic cells was determined by annexin V staining (E) (n=5). See also **Supp.** Figure 9. (F) Model: *S. aureus* interacts with an unknown primary receptor (1) that triggers production of NAADP, which in turn activates TPC1 to mediate lysosomal Ca^2+^ release and activation of Syt7 (2). Syt7-dependent lysosomal exocytosis (3) leads to release of ASM (4) and production of Cer on the plasma membrane (5) thereby presumably resulting in the recruitment of co-receptors [6,(84)] and rapid internalization of *S. aureus* by host cells (7). Bacteria, which enter the host cells via the rapid pathway, experience a different intracellular fate than bacteria that employed other pathways. **Statistics:** Mixed effects analysis (REML) and Tukey‘s multiple comparison test. Graphs represent mean ± SD. *p≤0.05.

Next, we tested whether internalization of the bacteria via the ASM-dependent pathway would affect phagosomal escape of *S. aureus*. We first confirmed that invasion of *S. aureus* is sensitive to ASM inhibitors and β-toxin treatment, despite reporter gene expression in the cell line (**Supp. Figure 5, E**). Then reporter cells either were left untreated or were pre-exposed to ASM inhibitors and were infected with *S. aureus* strain JE2. In addition, *S. aureus* Cowan I, a non-cytotoxic strain known to have low escape rates, was used for infection. The proportion of bacteria that escaped from the phagosome (**Figure 4, C, D**) as well as the proportion of escape events, which were additionally positive for cytosolic SM (**Supp. Figure 5, F**) were determined 1.5 h and 3 h p.i. (see also **Supp. Figure 7**, A-D). Whereas no differences for Lysenin-positive escape events were detected (**Supp. Figure 5, F**), a higher proportion of bacteria escaped from phagosomes on abrogation of ASM activity regardless of the inhibitor used (**Figure 4, C, D**). A similar increase was observed when we blocked lysosomal exocytosis by Vacuolin-1 ((**Figure 4, E**), suggesting that the lack of ASM delivery to the host cell surface during bacterial invasion affects phagosomal escape.

Since blocking of ASM-dependent uptake reduced invasion efficiency yet increased phagosomal escape rates, we hypothesized that the presence of SM on the plasma membrane affects downstream events during *S. aureus* infection. To address this, we pretreated host cells with the bacterial SMase β-toxin to block SM-dependent invasion (**Figure 5, A**; 2a, 2b) and either infected the cells in presence (**Figure 5, A;** 3a) or after removal of the toxin (3b). β-toxin pretreatment, resulted in enhanced phagosomal escape rates of *S. aureus* JE2, independent of the presence of β-toxin during infection (**Figure 5, B**; 4a, 4b). This was not observed for infections with the Cowan I strain, indicating that removal of SM by β-toxin did not affect general phagosomal membrane stability (**Supp. Figure 5, G**). Moreover, β-toxin treatment strongly reduced recruitment of Lysenin^W20A^, demonstrating effective removal of SM from membranes by β-toxin (**Figure 5, C** and **Supp. Figure 7, E-J**).

To investigate if removal of SM was important at the cell surface or within the phagosome, we infected the cells with transgenic *S. aureus* JE2 strain either collinearly overexpressing β-toxin and cyan-fluorescent reporter Cerulean or solely expressing Cerulean. We did not observe increased phagosomal escape upon overexpression of β-toxin (**Figure 5, B**; 4c), although Lysenin recruitment to phagosomal membranes was reduced to levels of β-toxin-pretreated samples (**Figure 5, C**). Moreover, we did not detect substantial differences in escape rates of *S. aureus* 6850, a strain naturally producing β-toxin (**Supp. Figure 5, H**), and its isogenic β-toxin mutant. Again, escape of *S. aureus* 6850 was enhanced upon pretreatment with β-toxin (**Supp. Figure 5, I**) accompanied by absence of SM from phagosomal membranes (**Supp. Figure 5, J**).

Phagosomal escape of *S. aureus* thus is independent of SM levels within the infected phagosomes but rather reflects the presence of SM and ASM at the plasma membrane during host cell entry.

We previously showed that *S. aureus* predominantly enters host cells via the rapid ASM- and Ca^2+^-dependent pathway early upon host cell contact (**Figure 1, C**; **Figure 2, F**; **Figure 3, D**), while other pathways are likely employed later during infection. Consequently, the proportion of pathways (ASM-dependent vs. ASM-independent) that is used for host cell entry relies on the infection time. To demonstrate that phagosomal escape is dependent on the pathway of cell entry, we infected reporter cells with *S. aureus* for either a 10 min (“early invaders”) or a 30 min infection pulse (representing “early and late invaders”) and at 3h p.i. we determined phagosomal escape rates (**Figure 5, D**) as well as the number of invaded bacteria (**Supp. Figure 5, K**). Independent of the multiplicity of infection (MOI) used, bacteria taken up within the first 10 min demonstrated significantly lower escape rates when compared to bacteria that were cocultured with host cells for 30 min.

This was corroborated by HeLa YFP-CWT infected with *S. aureus* expressing a fluorescent protein (e.g. mRFP) for 30 min (“early and late invaders”). After 20 min, we initiated a second infection pulse for 10 min (“early invaders”) with a *S. aureus* strain expressing another fluorescence protein (e.g. Cerulean) and determined phagosomal escape rates of both bacterial recombinants (**Figure 5, E**). Again, early invaders demonstrated lower escape rates when compared to the 30 min control. This was independent of the fluorescent marker expressed by the strain that was used for each infection pulse.

Taken together, we observed higher escape rates 3h p.i., whenever we blocked the rapid ASM-dependent invasion pathways. Consistently, escape rates of bacteria predominantly internalized via the ASM-dependent route (“early invaders”) were decreased (**Figure 5, F**). Hence, we concluded that the pathway of host cell entry directly affects phagosomal escape.

We next monitored phagosomal escape over 10 h in cells pretreated with ASM inhibitors (**Figure 6, A**, see **Supp. Figure 8**, for absolute bacteria numbers of individual replicates). ASM inhibition enhanced escape rates 3 h p.i. when compared to untreated controls. In contrast, 6 h p.i. escape rates were found decreased upon ASM inhibition.

Taken together, the efficiency of phagosomal escape of *S. aureus* is directly affected by the mode of cell entry, whereby the rapid SM/ASM-dependent pathway results in delayed phagosomal escape, while bacteria that entered host cells ASM-independently escape earlier during infection.

Since translocation to the host cytosol is a prerequisite for *S. aureus* replication in non-professional phagocytes, we recorded intracellular growth of *S. aureus.* In untreated controls replication started about 5 h p.i., and thus shortly after phagosomal escape (3-4 h p.i., **Figure 6, B**). By contrast, we did not observe significant bacterial replication in cells treated with ASM inhibitors (**Figure 6, B**).

We next treated HuLEC with β-toxin, stained the cells with BODIPY-FL-C12-SM and infected the cells with fluorescent *S. aureus*. Live cell imaging demonstrated that most untreated host cells died after 11 h p.i., whereas β-toxin-treated samples showed higher survival rates (**Figure 6, C**). This was also observed by cell death assays measuring i) membrane integrity (7-AAD staining, **Figure 6, D**), ii) apoptosis (Annexin V staining, **Figure 6, E**), iii) number of host cells that remained attached to the substratum (**Supp. Figure 9, A**) and iv) host cell lysis by LDH release (**Supp. Figure 9**, B) at 21 h p.i..

## Discussion

ASM has been previously reported to be implicated in the invasion of several viral, bacterial, and eukaryotic pathogens (7, 8, 13–17). However, the dynamics of ASM-mediated host cell entry, the underlying receptors as well as effects on post-invasion events are only poorly understood.

We here show that ASM is involved in the uptake of *S. aureus* by certain human cells within minutes after the pathogen contacts the host cell and this pathway is concurrent with ASM-independent uptake pathways. We demonstrate that the infection outcome of bacteria that enter the host cells via this rapid pathway is distinct from other entry mechanisms.

We observed a reduction of bacterial invasiveness upon treatment with different ASM inhibitors (**Figure 2, G**) or removal of the enzyme substrate SM from the plasma membrane (**Figure 2, N**) suggesting that ASM activity on the host cell surface is crucial for pathogen internalization. Since it remains elusive to which extent ASM processes SM on the plasma membrane during *S. aureus* invasion, one may speculate that ASM could also have functions other than SM metabolization during host cell entry of the pathogen. However, we did not detect a direct interaction between *S. aureus* and ASM in an *S. aureus*-host interactome screen (84).

Surprisingly, the uptake of *S. aureus* by ASM-deficient cells was only reduced relative to the wild type when cells were cultured in 1% but not in 10% FBS, a concentration commonly used in culture media (**Figure 2, I**). In ASM K.O.s cultured in 10% FBS, *S. aureus* invasion was only slightly affected by the ASM inhibitor ARC39, while ARC39-treated wildtype cells showed a stronger reduction (**Figure 2, J**). We concluded that reduced invasion in wildtype cells upon ARC39 treatment is mostly due to inhibition of ASM and consistently, the effect of ARC39 is strongly attenuated in ASM K.O.s. Hence, we postulate that other ASM-independent host cell pathways are upregulated in ASM-deficient cells cultured in 10% FBS, which might be caused by the strongly altered sphingolipid profile of these cells (**Figure 2, L, M**; **Supp. Figure 3**). Consistently, culturing ASM-deficient cells in 1% FBS resulted in less severe SM accumulation (**Figure 2, M**; **Supp. Figure 3, B** and D) and a reduced *S. aureus* invasion compared to wildtype cells (**Figure 2, I**).

For instance, *S. aureus* internalization by host cells is strongly restricted by caveolin-1, a protein associated with lipid membrane domains (78, 79). Accordingly, alterations in membrane composition might affect distribution of caveolin-1 on the plasma membrane, thereby fostering uptake of *S. aureus*. In consistency, previous studies reported dysfunction of caveolin-1-dependent endocytic processes in ASM-deficient cells (80, 81).

Release of ASM is caused by lysosomal exocytosis (6). Conventional methods for detection of lysosomal exocytosis often depend on the usage of anti-LAMP1 antibodies that are added to the cell culture medium for monitoring LAMP1 exposure on the plasma membrane (66). However, most antibodies are non-specifically bound by staphylococcal protein A (85) and hence, these protocols are not compatible with *S. aureus* infection studies. Therefore, we developed a novel approach to measure lysosomal exocytosis via a split luciferase-based assay that demonstrated lysosomal exocytosis during *S. aureus* infection (**Figure 2, A**-C). Consequently, depleting host cells from lysosomes that can undergo Ca^2+^-dependent exocytosis by ionomycin pretreatment, reduced invasion, even though ionomycin can influence general host cells Ca^2+^ homeostasis (86). Additionally, the lysosomal exocytosis inhibitor Vacuolin-1 resulted in drastically reduced internalization of bacteria (**Figure 2, D**), although the substance is known for side effects such as inhibition of PIKfyve thereby affecting the endo-lysosomal compartment (87). However, we confirmed the role of lysosomal exocytosis for *S. aureus* invasion of host cells by genetic ablation of Syt7, a key protein in lysosomal exocytosis (69).) It remains elusive, which other factors are involved in delivery of lysosomes to and fusion of lysosomes with the plasma membrane. Next to Syt7, there might be other proteins such as Synaptotagmin 1 with overlapping function that could compensate for the absence of Syt7 (88).

Elevation of cytosolic Ca^2+^ levels lead to recruitment of lysosomes to the cell surface (67). Whereas most studies implicated Ca^2+^ influx from the extracellular milieu (7, 16), we here demonstrate that *S. aureus* mainly triggers lysosomal Ca^2+^ mobilization as was shown by treatment with the inhibitor *trans*-Ned19 as well as a genetic K.O. of the endo-lysosomal Ca^2+^ channel TPC1 (**Figure 1, B** and E; **Supp. Figure 1, C**). Since fluorescent calcium reporters allow to monitor this process microscopically (89, 90), future experiments may visualize this process in more detail and contribute to our understanding of the underlying signaling mechanisms.

There are two TPCs (TPC1 and TPC2) that mediate lysosomal Ca^2+^ liberation (91), which have previously been associated with regulation of phagocytic processes (90) and host cell entry of several viruses such as Ebola virus (92), Middle East respiratory syndrome coronavirus [MERS-CoV, (93)] or SARS-CoV-II (94). In line with our study, absence of one of the two TPCs severely interfered with infection (92, 93).

TPC-dependent Ca^2+^ liberation was shown to induce exocytosis, for instance, during release of cytolytic granules from T cells (95) or during secretion of insulin (96). Our observations suggest that TPC1 is also involved in exocytosis of lysosomes, since exogenous addition of a bacterial SMase immediately before infection rescued the invasion defect in TPC1 and Syt7 K.O. cells (**Figure 2, P**) indicating that reduced invasion results from a decreased delivery of ASM to the host cell surface. Thus, NAADP-dependent Ca^2+^ liberation from lysosomal stores mediates Syt7-dependent fusion of lysosomes with the plasma membrane resulting in the release of ASM (**Figure 6, F**). While ionomycin treatment led to exocytosis of all “releasable” lysosomes, infection with *S. aureus* led to lysosomal exocytosis of only ∼30% of the ionomycin control (**Figure 2, C**), suggesting that *S. aureus* likely triggers local lysosomal exocytosis predominantly at host-bacteria contact sites, whereas ionomycin acts globally on all cells. A similar scenario has been observed for exocytosis of cytolytic granules in T lymphocytes, which differed between TPC/NAADP- and ionomycin-induction (95).

The conversion of SM to ceramide by ASM has been associated with the generation of ceramide-enriched platforms, lipid microdomains which facilitate the recruitment of certain protein receptors (97, 98), although the existence of such microdomains is still debated (99). We previously observed the interaction of *S. aureus* with several host cell surface receptors such as NRCAM or PTK7. The interaction of *S. aureus* with several host receptors was reduced upon blocking ASM-dependent internalization of *S. aureus* (84). It is thus tempting to speculate that accessory receptors identified in this study may be recruited to bacteria-host contact sites to facilitate rapid pathogen internalization.

Inhibition of CD38, a known NAADP producer (56, 57), did not influence uptake of staphylococci (**Supp. Figure 1, H**). By contrast, we detected a decreased invasion in cells deficient for the Toll-like receptor adaptor protein SARM1, an alternative NAADP producer [**Figure 1, D**; (59)]. However, the role of SARM1 in lysosomal exocytosis and ASM release requires further investigation. The impact of SARM1 on internalization of *S. aureus* was rather moderate, suggesting the existence of other NAADP-generating enzymes, such as the recently identified NADPH oxidases DUOX1, DUOX2, NOX1 and NOX2 (60).

One important characteristic of the ASM-dependent internalization pathway is its velocity - taking place within the initial ten minutes of an infection with *S. aureus*, during which treatment with small molecules (**Figure 3, D**), as well as genetic ablation of TPC1 (**Figure 1, C**) and Syt7 (**Figure 2, F**) resulted in reduction of intracellular bacteria. This was accompanied by the quick formation of *S. aureus*-containing phagosomes decorated with fluorescent SM analogs (**Figure 3, A**; **Supp. Figure 4**, **Supp. Video 1**, **Supp. Video 2**).

Neither treatment with inhibitors or enzymes nor genetic ablation blocking the rapid internalization pathway abolished the bacterial invasion of host cells completely. This supports our hypothesis that several concomitant host cell entry pathways exist, which is corroborated by a large number of host cell receptors interacting with *S. aureus* (13, 28, 31, 33–38). This pathway exists only in certain host cell types, since among the cell types, as is - for instance-illustrated by our experiments in which internalization of *S. aureus* was only sensitive to ASM inhibition in endothelial (HuLEC and HuVEC) and HeLa cells (**Figure 2, G**). The proportion of bacteria taken up by rapid internalization is dependent on the duration of infection and is higher in early (∼50-80% of all internalized bacteria 10 min p.i.) compared to late infection (∼15-50% of all internalized bacteria 30 min p.i.).

For simplicity we describe the pathway investigated in this study as “rapid” or “ASM-dependent”, however a clear cut between this and other pathways cannot be easily delineated. Additionally, internalization factors such as host cell receptors might possess overlapping function and could be involved in several pathways. In this context, ASM might simply accelerate certain internalization pathways, which could also take place in a slower fashion without involvement of ASM.

When we blocked the rapid internalization pathway by ASM inhibitors (**Figure 4, A, C, D**; **Supp. Figure 5, B**), Vacuolin-1 (**Figure 4, E**) or by removal of surface SM (**Figure 5, B**; **Supp. Figure 5, B, C**), we observed changes in both, maturation of bacteria-containing vesicles and phagosomal escape. A delayed maturation of phagosomes may affect acidification of the vesicles, a property that is sensed by *S. aureus* leading to expression of a different subset of virulence factors (100) and hence may alter intracellular pathogenicity.

The bacterial SMase β-toxin originates from *S. aureus*, however, most human pathogenic *S. aureus* strains including JE2, the strain we use in the present study, do not produce β-toxin, due to a genomic integration of a prophage (101). Phagosomal escape of *S. aureus* only was dependent on SM on the plasma membrane during invasion but not on the presence of SM within the phago-endosome (**Figure 5, A-C**; **Supp. Figure 5, I, J**). This not only corroborated that phagosomal escape of clinically relevant *S. aureus* strains is independent of the bacterial SMase β-toxin (42), but also demonstrated that the outcome of phagosomal escape is influenced by processes at the plasma membrane during cell entry. To exclude that inhibitor or bacterial SMase treatment caused changes in phagosomal escape by affecting cellular processes other than bacterial uptake, we infected untreated wildtype cells with *S. aureus* for 10 or 30 min, respectively, and monitored phagosomal escape. Bacteria that invaded within the first 10 minutes after contacting host cells (“early invaders”) are predominantly ASM-dependent (**Figure 3**) and demonstrated significantly less phagosomal escape (**Figure 5, D, E**). Consistently, blocking the ASM-dependent pathway led to increased proportions of ASM-independent invaders, which in turn resulted in higher escape rates (**Figure 4, C**-F; **Supp. Figure 5, G**). Intracellular *S. aureus* infections thus are directly influenced by the mode of internalization. This ultimately resulted in lower host cytotoxicity (**Figure 6, C**-E; **Supp. Figure 9**) and bacterial replication (**Figure 6, B**) when we blocked the ASM-dependent invasion.

An uptake-related intracellular fate has been suggested for *Mycobacterium bovis* (46), *Toxoplasma gondii* (48) or *Brucella abortus* (49). However, in these studies the host cells were either depleted from cholesterol, which causes a variety of side effects (47), or internalization was artificially induced by chemicals or ectopic expression of receptors, respectively.

Taken together, we here describe a rapid internalization pathway for *S. aureus* into human epithelial and endothelial cells. Bacteria that enter the host cells via this infection route face a distinct intracellular fate. Here we show, to our knowledge, for the first time that host cells within the same population can be invaded by a pathogen via multiple routes and that this results in a differential infection outcome. Rapid internalization may prove beneficial for pathogens in an *in vivo* setting since internalization by host cells shortens the exposure to the innate immune system within a host. Interestingly, the absence of ASM in *Smpd1*^-/-^ mice enhanced the potency of antibiotics to clear *S. aureus* sepsis (102). It is tempting to speculate that reduced host cell invasion in absence of ASM caused a prolonged exposure of the pathogen to antibiotics. Several FIASMAs are already approved drugs (103), and may have future use in infection treatment.

## Acknowledgements

This work was supported by the Deutsche Forschungsgemeinschaft (DFG; https://www.dfg.de) within the research training group RTG2581 (to M.F., F.Schu., B.K.) and CRC1583 (to M.F.). C.K. and C.A. are grateful to the DFG AR 376/22-1. M.R. was supported by funds of the Bavarian State Ministry of Science and the Arts and the University of Würzburg to the Graduate School of Life Sciences (GSLS), University of Würzburg. The DFG funded the Leica TCS SP5 CLSM under project code 116162193 and the BD FACSAriaIII cell sorter under project code 206080318.

We thank Sibylle Schneider-Schaulies and Thomas Rudel for valuable discussions and critically reading the manuscript, and Nadine Vollmuth as well as Christian Stigloher for scientific advice and intense discussions. We further thank Daniel Herrmann for assistance with HPLC-MS/MS measurements of sphingolipids. We are indebted to Kathrin Stelzner for conducting FACS of cell lines and generation of *S. aureus* strains. We also are grateful to Norbert Klugbauer (Freiburg, Germany) for providing TPC1/TPC2 knock-out cells. pmRFP-LC3 was a gift from Tamotsu Yoshimori (Addgene plasmid # 21075; http://n2t.net/addgene:21075; RRID:Addgene_21075, (104)). pLVTHM (105), pMD2.G and psPAX2 (unpublished) were a gift from Didier Trono (Addgene plasmids # 12247, 12259 and 12260, respectively). pLX304-Flag-APEX2-NES was a gift from Alice Ting (Addgene plasmid # 92158, (106)) and pSpCas9(BB)-p2A-GFP (PX458) (74) was a gift from Feng Zhang (Addgene plasmid # 48138; http://n2t.net/addgene:48138). Figures were created in BioRender.

## Author Contributions

M.R., M.F. conceptualized the work, designed the methodology and wrote the original draft manuscript. M.F., B.K. acquired funding. M.R. F.S., K.U., M.P., A.M., N.K., J.W., A.I., C.K., K.P., F. Schu performed the experiments; M.R., M.F. supervised the experiments. M.R., C.A., M.F. analyzed the data; all authors edited and revised the manuscript.

## Materials and Methods

### Cell culture

HeLa (ATCC CCL-2^TM^), 16HBE14o-(kindly provided by Prof. Jan-Peter Hildebrandt, University of Greifswald) and EA.hy 926 (107) were cultivated in RPMI+GlutaMAX^TM^ medium (Gibco^TM^, Cat. No. 72400054) supplemented with 10% (v/v) heat-inactivated (56°C at 30 min) fetal bovine serum (FBS, Sigma Aldrich, Cat. No. F7524) and 1 mM sodium pyruvate (Gibco^TM^, Cat.No. 11360088).

HuLEC-5a (ATCC CRL-3244^TM^) and HuVEC (Gibco^TM^, Cat. No.C01510C) were cultured in MCDB131 medium (Gibco^TM^, Cat. No. 10372019) supplemented with microvascular growth supplement (Gibco^TM^, Cat. No. S00525), 2 mM GlutaMAX^TM^ (Gibco^TM^, 35050061), 5 % (v/v) heat-inactivated (56°C at 30 min) FBS, 2.76 µM hydrocortisone (Sigma Aldrich, Cat. No. H0888), 0.01 ng/ml human epidermal growth factor (Pep Rotech, AF-100-15) and 1x Penicillin-Streptomycin (Gibco^TM^, Cat. No. 15140122).

#### HeLa

HuLEC and HuVEC were detached by StemPro^TM^ Accutase^TM^ (Gibco^TM^, Cat. No. A1110501) and seeded at the indicated density two days prior to the experiment, whereas HeLa, 16HBE14o- and EA.hy 926 were detached with TrypLE^TM^ (Gibco^TM^, Cat. No. 12604013) and seeded one day prior to the experiment.

When experiments were to be conducted with only 1% FBS in the culture medium, the cells initially were cultured in media containing 10% FBS, were shifted to media containing 2% FBS for up to three days and eventually were cultivated in media with 1% FBS one day prior to the experiment.

### Generation of HeLa KO cell lines

Generation of K.O. cell lines is based on a previous protocol (74). sgRNAs (**Table 1**) were designed with the CHOPCHOP online tool (chopchop.cbu.uib.no) and synthesized as primer pairs (Sigma Aldrich). After phosphorylation with T4 nucleotide kinase (ThermoFisher, Cat. No. ER0031), the sgRNAs were cloned into the BbsI sites of pSpCas9(BB)-p2A-GFP (AddGene #48138, (74)) via forced ligation by BbsI (ThermoFisher, Cat. No. ER1012) and T4 Nulceotide ligase (ThermoFisher, Cat. No. EL0011). The constructs were then transformed in *E. coli* DH5α (ThermoFisher, Cat. No EC0112) and subsequently sequence verified.

**Table 1.**
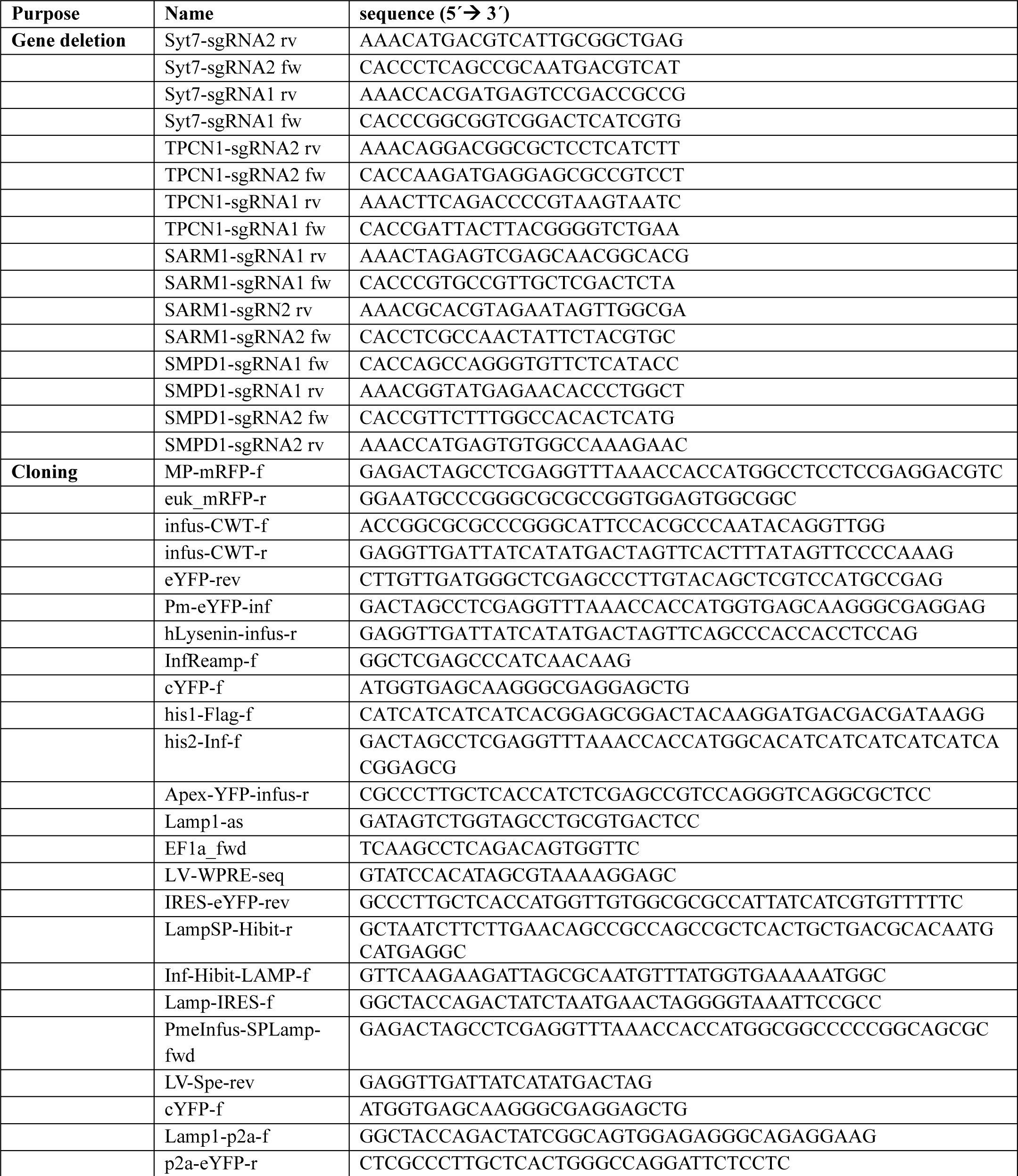
Oligonucleotides used in this study.

HeLa cells were transfected with 1 μg each of two distinct sgRNA constructs using JetPrime (Polyplus) following the manufacturer’s instructions. After 36-48h, cells were sorted for strong GFP expression in a FACS Aria III (BD Bioscience).To isolate ASM K.O. cell lines, we included a second sorting step. Therefore, the cells were incubated with 10 µM FRET probe (73) for 2 h at 37°C/5% CO_2_ and sorted for cells negative in FITC emission, which correlates with the cellular ASM activity. The resulting cell pools were cultivated and tested for loss of expression of the respective sgRNA target by Western Blotting or ASM activity assays.

The following antibodies were used: mouse anti-PTK7 monoclonal IgG1 (bio-techne, Cat. No. MAB4499) with a horse radish peroxidase (HRP)-conjugated anti-mouse antibody (SantaCruz, Cat. No. sc-516102) and rabbit anti-TPC1 polyclonal (Thermo Fisher, Cat. No. PA5-41048) or rabbit anti-SARM1 monoclonal antibody (Cell Signaling Technology, Cat. No. 13022) with an HRP-conjugated anti-rabbit antibody (Biozol, Cat. No. 111-035-144).

### Generation of reporter constructs

*pLV-mRFP-CWT.* mRFP and the cell wall-targeting domain of lysostaphin CWT were amplified using primers MP-mRFP-f and euk_mRFP-r, as well as infus-CWT-f and infus-CWT-r, respectively. As PCR templates we used pmRFP-LC3 (PMID 17534139) and pLV-YFP-CWT (108). The resulting plasmid was termed pLV-mRFP-CWT. The PCR products were subsequently cloned into pLVTHM (105) restricted with PmeI and SpeI by InFusion cloning following the manufacturer’s recommendations.

*pLV-YFP-Lysenin.* The sequence of the sphingomyelin-binding but not pore-forming earthworm toxin mutant Lysenin^W20A^ (83) was synthesized by GeneArt (ThermoFisher) adapted to human codon usage and with a 5’ flanking region encoding a glycine-rich linker sequence. The synthesized fragment served as template for subsequent PCRs.

Lysenin^W20^ was amplified using oligonucleotides InfReamp-f and hLysenin-infus-r (**Table 1**).

YFP was amplified with Pm-eYFP-inf and eYFP-rev from template pLVTHM-YFP-CWT (108). Both PCR products were joined with pLVTHM vector restricted with PmeI and SpeI by InFusion cloning following the manuacturer’s recommendations, thereby creating pLV-YFP-Lysenin.

Constructs were transformed into *E. coli* DH5α and sequences were validated. Plasmid preparations were performed using standard laboratory procedures. VSV-G-pseudotyped Lentiviral particles were generated by transfecting with each lentiviral vector as well as the plasmids pMD2.G (VSV G) and psPAX2 following the protocol of (105) and target cells were transduced as described previously (108).

### Bacteria culture

*S. aureus* liquid cultures were either grown in 37 g/L brain heart infusion (BHI) medium (Thermo Fisher, Cat. No. 237300) or in 30 g/l TSB medium (Sigma Aldrich, Cat. No. T8907). *E. coli* liquid cultures were grown in LB medium [10 g/l tryptone/peptone (Roth, Cat. No. 8952.4,) 5g/l yeast extract (Roth, Cat. No 2363.2), 10 g/l NaCl (Sigma Aldrich, Cat. No. S5886)].

For agar plates, 15 g/L agar (Otto Norwald, Cat. No. 257353), was added to either TSB (*S. aureus*) or LB (E. coli).

Media and plates were supplemented with appropriate antibiotics [5 µg/ml erythromycin (Sigma Aldrich, Cat. No. E5389), 10 µg/ml chloramphenicol (Roth, Cat. No. 3886.2), 100 ng/ml Carbenicillin (Roth, Cat. No. 6344.2)].

Strains used in this study are listed in **Table 2**.

**Table 2.** Bacterial Strains used in this study.

| Strain | Remarks | Source |
| --- | --- | --- |
| *S. aureus* 6850 |  | (109) |
| *S. aureus* 6850 $\Delta hlb$ | deficient in β-toxin production | (42) |
| *S. aureus* Cowan I | NCTC 8530, isolated from septic arthritis, agr dysfunction, low expression of toxins and proteases | ATCC 12598 |
| *S. aureus* JE2 | MRSA, USA300 | (110) |
| *S. aureus* JE2 $\Delta hla$ | deficient in α-toxin production | (111) |
| *S. aureus* JE2 $\Delta hlb$ | deficient in β-toxin production | (11) |
| *S. aureus* JE2 SarAP1 Cerulean | Constitutive expression of the fluorescence protein Cerulean | (50) |
| *S. aureus* JE2 SarAP1 mRFP | Constitutive expression of the fluorescence protein mRFP | (45) |
| *S. aureus* JE2 pCer | Anhydrous tetracycline (AHT) inducible expression of Cerulean, construct published in (112) and was transduced into *S. aureus* JE2 | This work |
| *S. aureus* JE2 pCer+hlb | Anhydrous tetracycline (AHT) inducible expression of Cerulean and β-toxin, construct published in (112) and was transduced into *S. aureus* JE2 | This work |
| *S. aureus* Cowan I SarAP1 Cerulean | Constitutive expression of the fluorescence protein Cerulean | This work |
| *S. aureus* 6850 SarAP1 Cerulean | Constitutive expression of the fluorescence protein Cerulean | (45) |
| *S. aureus* 6850 $\Delta hlb$ SarAP1<br>Cerulean | Constitutive expression of the fluorescence protein<br>Cerulean | This work |

### *S. aureus* growth curves

*S. aureus* overnight cultures were grown in BHI medium, and 1 ml was centrifuged at 14,000 for 1 min. Bacteria were washed thrice with DPBS and either resuspended in BHI (for amitriptyline and ARC39 treatment) or infection medium [MCDB131 medium (Gibco^TM^, Cat. No. 10372019) complemented with 2 mM GlutaMAX^TM^ (Gibco^TM^, 35050061) and 10 % (v/v) heat-inactivated (56°C at 30 min) FBS, for ionomycin treatment). Bacteria suspensions were diluted to OD_600_=0.1 in the respective medium. Then, 400 µL per well of the suspension as well as respective blanks were transferred into a 48 well plate and OD_600_ was determined every 16 min in Tecan mPlex200 microplate reader.

### *S. aureus* infections

Host cells were seeded in 6 well plates (2×10^5^ cells per well), 12 well plates (1×10^5^ cells per well) or 24 well plates (0.5×10^5^ cell per well) either one day (HeLa, EA.hy 926, 16HBE14o^-^) or two days (HuLEC, HuVEC) prior to the experiment. Host cells were washed thrice with DPBS and infection medium or Ca^2+^-free infection medium [DMEM w/o calcium (Gibco^TM^, Cat. No. 21068028) complemented with 10 % (v/v) heat-inactivated (56°C at 30 min) FBS and 200 µM BAPTA (Merck Millipore, 196418)] with or without 1.8 mM CaCl_2_ (Roth, Cat. No. 5239.1) was added. If indicated, cells were pretreated with compounds prior to infection (for details about treatments see **Table 1**). For experiments involving blocking of receptors by antibodies, respective solvent controls were implemented (final concentrations in infection medium: 0.002% (w/v) NaN_3_ (Sigma Aldrich, Cat. No. S2002) 5% glycerol (Roth, Cat. No 3783.2) in 10% DPBS for 30 ng/ml anti-NRCAM antibody as well as 0.01 % (w/v) NaN_3_ for 50 ng/ml anti-CD73 and 50 ng/ml anti-MELTF antibodies.

An *S. aureus* overnight culture grown in BHI containing the appropriate antibiotics was diluted to an OD_600_=0.4 in the same medium. When inducible expression of genes was required for the strains *S. aureus* JE2 p*Cer* and *S. aureus* JE2 p*Cer*+*hlb,* 200 ng/ml anhydrous tetracycline (AHT, AcrosOrganics, 233131000) was added. The culture was grown to an OD_600_= 0.6-1.0 and 1 ml bacterial suspension was harvested by centrifugation and washed twice with Dulbecco’s Phosphate Buffered Saline (DPBS, Gibco^TM^, Cat. No 14190169). The bacteria were resuspended in infection medium (or Ca^2+^-free infection medium when infection was performed in absence of extracellular Ca^2+^). The number of bacteria per ml in the suspension was determined with a Thoma counting chamber and the MOI was determined. If not indicated otherwise, an MOI=10 was used for infections.

The infection was synchronized by centrifugation 800xg/8 min/RT (end of the centrifugation: t=0). To determine bacterial invasion, the infection was stopped after 30 min (unless indicated otherwise) by removing extracellular bacteria with 20 µg/ml Lysostaphin (AMBI, Cat. No. AMBICINL) in infection medium for 30 min. Then, host cells were washed thrice with DPBS, lysed by addition of 1 ml/well (12 well plate) or 0.5 ml/well (24 well plate) Millipore water and the number of bacteria in lysates was determined by plating serial dilutions (10^-1^, 10^-2^, 10^-3^) on TSB agar plates. Plates were incubated overnight at 37 °C and colony forming units (CFU) were enumerated. To determine invasion efficiency, the number of bacteria determined in tested samples were normalized to untreated controls (set to 100%). For measuring invasion dynamics, the number of bacteria was either normalized to the 30 min time point of untreated controls or to the corresponding time points of untreated controls.

If later time points in infection were investigated, infection medium containing 2 µg/ml Lysostaphin was added to the cells until the indicated time.

For testing bacterial susceptibility to inhibitors (“survival”) or adherence, Lysostaphin treatment was omitted, and host cells were immediately lysed by addition of Millipore water or washed five times with DPBS and subsequently lysed with Millipore water, respectively. The number of bacteria in lysates was determined by CFU counting (dilutions 10^-2^, 10^-3^, 10^-4^) on TSB agar plates.

### Phagosomal escape assays

For phagosomal escape assays with amitriptyline, ARC39, PCK310 and β-toxin, HeLa cells expressing RFP-CWT and Lysenin^W20A^-YFP were infected with the indicated *S. aureus* strain as described in the previous paragraph. Infections with *S. aureus* JE2 p*Cer* and *S. aureus* JE2 p*Cer*+*hlb* were carried out in presence of 200 ng/ml AHT.

For phagosomal escape assays, HeLa cells expressing RFP-CWT and Lysenin^W20A^-YFP were infected for 10 min (early invaders) or 30 min (early and late invaders) at the indicated MOIs.

For the phagosomal escape assays with two *S. aureus* JE2 strains expressing different fluorescence proteins, HeLa cells expressing YFP-CWT were infected with an MOI=5 of an initial *S. aureus* JE2 strain (*S. aureus* JE2 SarAP1 Cerulean or *S. aureus* JE2 SarAP1 mRFP). After 12 min of infection, a second infection pulse was initiated by adding the second *S. aureus* JE2 strain (expressing the complementary fluorescence protein, *S. aureus* JE2 SarAP1 mRFP or *S. aureus* JE2 SarAP1 Cerulean, respectively) and centrifuging the bacteria on the host cells. To exclude that the type of fluorescence protein expressed by the *S. aureus* strains contributes to the experimental outcome, either strain was used for the first and second infection pulse (each combination: n=4).

### Phagosomal maturation assays

To monitor phagosomal maturation, HeLa cells expressing mCherry-Rab5 and YFP-Rab7 or YFP-Rab7 and RFP-CWT were infected with *S. aureus* JE2 SarAP1 Cerulean.

After 3 h p.i. (unless indicated otherwise), cells were washed thrice with DPBS and fixed with 0.2% glutaraldehyde (Sigma Aldrich, Cat. No. 10333) / 4% paraformaldehyde in PBS (Morphisto, Cat. No 11762.01000) for 30 min at RT. Then, cells were washed thrice with DPBS, stained with 5µg/ml Hoechst 34580 (Thermo Fisher, Cat. No. H21486) and mounted in Mowiol [24g glycerol (Roth, Cat. No. 3783.2), 9.6 g Mowiol^®^4-88 (Roth, Cat. No 0713.2), 48 ml 0.2 M TRIS-HCl pH 8.5 (Sigma Aldrich, Cat. No T1503), 24 ml Millipore water].

Samples were imaged with a Leica TCS SP5 confocal microscope (Wetzlar, Germany; Software Leica LAS AF Version 2.7.3.9723) with a 40x immersion oil objective (NA1.3) and a resolution of 1024×1024 pixels [Cerulean (Ex. 458 nm/Em. 460-520), YFP (Ex. 514 nm/Em. 520-570nm), mRFP/mCherry (Ex. 561 nm/Em.571-635nm) and Hoechst 34580 (Ex. 405 nm/Em.410-460nm). At least 10 fields of view per sample were recorded.

Image analysis was performed in Fiji (113) using a previously described macro (112) that identifies and extracts individual bacteria from images as regions of interest (ROIs, see **Supp. Figure 7**). Subsequently, fluorescent intensity in every channel is measured to determine recruitment of fluorescence reporters to the bacteria. Results were exported as text files and proportion of bacteria that recruited individual reporters was determined with the Flowing 2 software (Turku Bioscience Center). Phagosomal escape rates were determined as the proportion of CWT-positive bacteria of the total number of intracellular bacteria. Similarly, the proportion of Lysenin^W20A^-positive escape events, the ratio of Lysenin^W20A^-/CWT-positive events and all CWT-positive events was calculated.

To measure the proportion of bacteria that were associated with Rab5 and/or Rab7, the proportion of Rab5- and/or Rab7-positive membranes around bacteria of all intracellular bacteria was determined. For experiments obtained with HeLa cells expressing YFP-Rab7 and RFP-CWT, bacteria that acquired RFP-CWT were removed from the dataset. Subsequently, the proportion of Rab7-positive bacteria in relation to all CWT-negative intracellular bacteria was determined.

### Live cell imaging

The indicated host cells were seeded one (HeLa) or two days (HuLEC) prior to the experiments in µ-slide 8 well live cell chambers (ibidi, Cat. No 80826-90) with a density of 0.375×10^5^ cells per well.

For monitoring host cell entry, cells were washed thrice with DPBS and were then incubated for 90 min in infection medium with 1 µM BODIPY-FL-C_12_-sphingomyelin (Thermo Fisher Cat. No. D7711), 10 µM BODIPY-FL-C_12_-ceramide (Santa Cruz, Cat.No. sc-503923) or 10 µM visible-range FRET probe (73). Pretreatment with the FRET probe was performed in infection medium containing 1 % (v/v) FBS. Then, cells were washed thrice with DPBS and imaging medium [RPMI 1640 w/o phenol red (Gibco^TM^, Cat. No. 11835030) containing 10% (v/v) heat-inactivated (56°C/30 min) FBS (Sigma Aldrich, Cat. No. F7524) and 30 mM HEPES (Gibco^TM^, Cat.No. 15630080)] applied. Samples stained with the FRET probe were imaged in imaging medium containing 1% FBS and in presence of 10 µM FRET probe. Samples were infected with *S. aureus* JE2 SarAP1 Cerulean (FRET probe samples) or *S. aureus* JE2 SarAP1 mRFP (BODIPY-FL-C12-sphingomylein/-ceramide) at an MOI=50 without synchronization by centrifugation. Infection was monitored with a Leica TCS SP5 confocal microscope (Wetzlar, Germany) in intervals of 1 min by recording BODIPY-FL (Ex. 496 nm/Em. 500-535 nm), mRFP (Ex. 561 nm/Em.571) FITC (Ex. 488 nm/Em. 500-560), Cerulean (Ex. 458 nm/Em. 460-520), BODIPY-TR (Ex. 594 nm/Em. 610-680) and FRET (Ex.488 nm/ Em:610-680) channels. Cells were recorded with a 40x immersion oil objective and a resolution of 2048×2048 pixels. For samples stained with the FRET probe, 20 µg/ml Lysostaphin was added 40 min p.i. to remove extracellular bacteria.

For quantification of bacteria that associate with BODIPY-FL-C_12_-sphingomyelin/-ceramide, individual bacteria in single frames were identified, extracted as ROIs and BODIPY-FL fluorescence in ROIs was measured in Fiji (113) as described before for phagosomal escape assays. Results were extracted as text files and the proportion of bacteria associating with BODIPY-FL was determined with Flowing2 (Turku Bioscience Center).

For monitoring intracellular *S. aureus* infection, host cells were pretreated and infected with *S. aureus* JE2 SarAP1 Cerulean (HeLa RFP-CWT/Lysenin^W20A^-YFP) or *S. aureus* JE2 SarAP1 mRFP (HuLEC) as described in “*S. aureus* infection”. HuLEC additionally were treated with 1 µM BODIPY-FL-C_12_-sphingomyelin in infection medium for 90 min and washed thrice with DPBS before the indicated treatment was applied. After extracellular bacteria were removed with 20 µg/ml lysostaphin for 30 min, imaging medium containing 2 µg/ml lysostaphin was applied and infections were monitored in intervals of 5 min (HeLa RFP-CWT) or 20 min (HuLEC). Phagosomal escape was evaluated in Fiji as described above. To determine intracellular replication, the number of bacteria was determined for each individual frame as described before (112). The number of bacteria at each time point was then normalized to the number of bacteria detected in the first frame to calculate the relative replication.

### ASM and NSM activity assays

Thin layer chromatography (TLC)-based ASM/NSM activity assays were adapted from previously published protocols (72). To determine cellular ASM activity, the indicated cell lines were seeded in 24 well plates with a density of 1×10^5^ cell per well one day (HeLa, EA.hy 926, 16HBE14o^-^) or two days (HuLEC, HuVEC) prior to the experiment. Cells were either treated with 20 µM amitriptyline or with 10 µM ARC39 for 22 h (fresh inhibitor was applied after 18h) in infection medium. Then, cells were washed thrice and lysed by addition of 100 µL per well ASM lysis buffer [250 mM NaOAc pH 5 (Roth, Cat. No 6773.2), 0.1% Nonidet^®^ P40 substitute (AppliChem. Cat. No. A1694,0250), 1.3 mM EDTA 1.3 mM (Roth, Cat. No. 8040.2), 1x protease inhibitor cocktail (Sigma Aldrich, 11873580001)] for 15 min/4°C. Cells were scraped from the substratum and protein concentration in the resulting lysates was measured with a Pierce^TM^ bicinchoninic acid (BCA) assay kit (Thermo Fisher, Cat. No. 23227). 1 µg of protein was incubated with 100 µL ASM lysate assay buffer [200 mM NaOAc pH 5 (Roth, Cat. No 6773.2), 0.02 % Nonidet^®^ P40 substitute (AppliChem. Cat. No. A1694,0250), 500 mM NaCl (VWR, Cat. No. 27810.364)] containing 0.58 µM BODIPY-FL-C_12_-Sphingomyelin (Thermo Fisher, Cat. No. D7711) for 4h at 37°C and 300 rpm.

For measuring bacterial SMase/NSM activity in *S. aureus* cultures, an overnight culture of the indicated strain was grown in BHI medium, centrifuged 14.000xg and the supernatant was sterile filtered with 0.2 µm filter. 100 µL of sterile supernatant was incubated with 100 µL NSM assay buffer [200 mM HEPES, pH 7.0 (Roth, Cat. No. 6763.3), 200 mM MgCl_2_ (Roth, Cat.No. 2189.2), 0.05% Nonidet P-40 (AppliChem. Cat. No. A1694,0250)] for 4h/37°C/300 rpm.

The reactions were stopped by the addition of 2:1 CHCl_3_:MeOH (Roth, Cat. No. 3313.2 and Cat. No. 8388.6). Samples were vortexed, centrifuged at 13,000 x g for 3 min and 50-100 µL of the lower organic phase were transferred to a fresh tube. Samples were completely evaporated using a SpeedVac 5301 concentrator (Eppendorf), resuspended in 10 µL 2:1 CHCl_3_/MeOH and spotted in 2.5 µL aliquots on a TLC plate (Alugram, Xtra Sil G/UV254, 0.2 mm/silica gel 60; VWR, Cat. No. 552-1006).

Plates were developed using 80:20 CHCl_3_:MeOH and subsequently scanned with a Typhoon 9200 Scanner (Amersham). For quantification, intensities of the lower (SM) and the upper bands (Cer) were measured in Fiji (113) and activity was determined based on the reaction time, protein amount and SM/Cer ratios.

For the flow cytometry based read out, cells seeded in a density of 0.5×10^5^ cells per well in a 24 well plate either one day (HeLa, EA.hy 926, 16HBE14o^-^) or two days (HuLEC, HuVEC) prior to the experiment. Cells were treated with ARC39 and amitriptyline as described above. Then, cells were washed thrice with DPBS and incubated with 10 µM FRET probe (73) for 2h in presence of the inhibitors. Subsequently, cells were washed thrice, detached with TrypLE^TM^ (Gibco^TM^, Cat. No. 12604013) and resuspended with 2% (v/v) FBS in DPBS. Samples were analyzed for FITC (Ex. 488 nm/Em. band pass 530/30 nm) and BODIPY-TR (Ex.: 561nm/ Em. band pass 695/40nm) fluorescence with Attune NxT flow cytometer (Thermo Fisher, Attune Cytometric Software v5.2.0). Cell populations were analyzed for FITC and BODIPY-TR mean fluorescence in Flowing 2 (Turku Bioscience Center) and FITC vs. BODIPY-TR ratios were calculated to determine arbitrary probe conversion.

### Determination of β-toxin activity on living cells

HeLa cells were seeded in a 24 well plate with a density of 0.5×10^5^ cells per well. Cells were incubated with 1 µM BODIPY-FL-C_12_-SM for 24h. Then, samples were washed thrice with DPBS and treated with 100 ng/ml β-toxin in infection medium or left untreated for 75 min. Subsequently, cells were washed thrice with DPBS, lysed with 200 µL MeOH and detached with a cell scraper. 400 µL CHCl_3_ and samples were centrifuged for 5 min/25.000xg to remove cell debris. 100 µL of the sample were completely evaporated using a SpeedVac 5301 concentrator (Eppendorf), resuspended in 10 µL 2:1 CHCl_3_/MeOH and spotted in 2.5 µL aliquots on a TLC plate (Alugram, Xtra Sil G/UV254, 0.2 mm/silica gel 60; VWR, Cat. No. 552-1006). TLC was developed with 80:20 CHCl_3_:MeOH and scanned with a Typhoon RGB scanner (Amersham). Fluorescence intensities were evaluated in Fiji and ratios ceramide vs. SM were calculated.

### Construction of HiBit-SP-LAMP1-p2A-eYFP

To visualize the transient surface exposure of lysosomal membranes on the plasma membrane of epithelial cells we made use of a split NanoLuc luciferase. We inserted a region encoding the 11 amino acid residues encompassing HiBiT peptide of the split NanoLuc (Promega) between the signal peptide and the N-terminus of mature LAMP1.

For that purpose, we amplified a 144 bp fragment encoding the signal peptide of LAMP1 (SP_LAMP1_, PmeInfus-SPLamp-fwd & LAMP1SP-HiBiT-r), the 1.2 kb coding region of mature LAMP1 (mLAMP1, Inf-HiBiT-LAMP-f & LAMP1-as) and eYFP (cYFP-f & LV-Spe-rev) via PCR using the indicated primer pairs respectively and using LAMP1-YFP containing plasmid as DNA template (41). A p2A peptide encoding sequence was amplified with oligonucleotides Lamp1-p2a-f and p2a-eYFP-r using pspCas9 (74) as a template. As vector we used the PmeI/SpeI-restricted pLVTHM (105).

The resulting DNA Fragments contained overlapping ends of at least 16 bp, which were assembled by an InFusion enzyme mix (Takara Biotech) according to the manufacturer ‘s instructions. The resulting plasmid was transformed into chemically competent *E. coli* DH5α and plated on LB agar containing 100 μg/ml ampicillin. Colonies were inoculated, and plasmid were prepared from the resulting overnight cultures using the Qiagen Plasmid Mini kit. Sequences were determined using Sanger sequencing (Microsynth) thereby establishing the correct assembly of the vector. Pseudotyped lentivirus particles were generated in HEK293 cells and transduction of HeLa cells was conducted following a published protocol (105).

eYFP production in transgenic cells here served as reporter for transfection of the cells and production of the SP-HiBiT-LAMP1 proportion of the fusion protein. eYFP-positive cells were FACS-sorted for medium eYFP expression levels and the resulting cell pool was named HeLa SP_LAMP_1-HiBit-LAMP1-p2A-EYFP.

### Determination of externalization of lysosomal membranes using split NanoLuc biolumin-escence

For measurement of LAMP1 externalization, 4×10^4^ per well of HeLa SP_LAMP_1-HiBit-LAMP1-p2A-EYFP, hereafter named HeLa HiBiT-LAMP1, were seeded in white flat-bottom 96 well plates (Nunc™ MicroWell™ 96 Wells, Nunclon Delta-treated, Cat. No. 136101) and grown overnight yielding 8×10^4^ cells at the next day. Bioluminescence was detected using the Promega Nano-Glo HiBiT Extracellular Detection Kit (Promega # N2420).

For infection with *S. aureus*, bacteria from an overnight culture were diluted to an OD600 of approximately 0.5 into 10 ml TSB and grown for 1 h. In case a preincubation with inhibitors was required (e.g. as for vacuolin-1), cells were washed twice with DPBS, and inhibitors were added in indicated concentrations in a final volume of 200 µl per well.

Extracellular detection reagent was prepared by mixing 4 ml extracellular buffer, 20 µl LgBit protein and 40 µl bioluminescent substrate [Promega, Cat. No. N2420]. 100 µl extracellular detection reagent was added per well and luminescence was measured five times in 2 min intervals on a TECAN Infinite Pro 200 plate reader (37 °C, attenuation: OD1, integration time 250 ms). Then, inhibitors or bacteria were added accordingly, and luminescence measurements were continued in 2 min intervals using the settings above. The relative luminescence units (RLUs) determined in samples infected with *S. aureus* were scaled to the untreated control (set to 0%) and cells treated with 1 µM ionomycin (set to 100 %). The results describe the LAMP1 externalization as proportion of all lysosomes that can be released from host cells by ionomycin treatment.

### Determination of cellular sphingolipid profiles via HPLC-MS/MS

HeLa wildtype or ASM K.O. cells were seeded with a density of 3.5×10^6^ cells per well in a 6-well plate. wildtype cells were treated as described in **Table 3** with 10 µM ARC39 or 20 µM amitriptyline. Then, cells were washed thrice with DPBS, lysed with 250 µL cold MeOH and detached with a cell scraper on ice. Wells were rinsed with an additional 250 µL MeOH to ensure entire sample collection.

**Table 3.** Host cell treatment with various compounds.

| compound | Supplier/Cat.No. | preincubation time | removal prior to infection? |
| --- | --- | --- | --- |
| 2-APB<br>(2-Aminoethoxydiphenylborate) | Sigma Aldrich/<br>D9754 | 75 min | removed prior to infection |
| 8-Bromo-cADPR<br>(8-Bromo-cyclic ADP ribose) | Biolog/065-005 | 75 min | present during infection |
| amitriptyline | Sigma Aldrich/<br>A8404 | 75 min | present during infection |
| ARC39 | (71) | 18 h, add fresh inhibitor for another 4h or as indicated | present during infection |
| BAPTA-AM | Merck Millipore/<br>196419; (51) | 25 min | removed prior to infection |
| BODIPY-FL™-C12-Ceramide | Santa Cruz/<br>sc-503923 | 90 min | present during infection |
| BODIPY-FL™-C12-Sphingomyelin | ThermoFisher/D7711 | 90 min | present during infection |
| CD38i/78c | Sigma Aldrich/<br>5387630001 | 75 min | present during infection |
| Ionomycin | Sigma Aldrich/ I0634 | 75 min | present during infection |
| PCK310 | To be published elsewhere, patent pending | 4h or as indicated | present during infection |
| *trans*-Ned19 | Sigma Aldrich/ SML2830; (55) | 75 min | present during infection |
| Vacuolin-1 | Biozol/ MCE-HY-118630 | 75 min | present during infection |
| β-toxin | Sigma Aldrich/ S8633 | 75 min | removed prior to infection if not indicated otherwise |

Cell suspensions were then subjected to lipid extraction using 1 mL MeOH/CHCl_3_ (2:1, v:v) as described before (114). The extraction solvent contained C17 ceramide (C17 Cer) and d_31_-C16 sphingomyelin (d_31_-C16 SM) (both Avanti Polar Lipids, Alabaster, USA) as internal standards. Chromatographic separations were achieved on a 1290 Infinity II HPLC (Agilent Technologies, Waldbronn, Germany) equipped with a Poroshell 120 EC-C8 column (3.0 × 150 mm, 2.7 µm; Agilent Technologies). MS/MS analyses were carried out using a 6495C triple-quadrupole mass spectrometer (Agilent Technologies) operating in the positive electrospray ionization mode (ESI+). Cer and SM were quantified by multiple reaction monitoring (qualifier product ions in parentheses): [M-H_2_O+H]^+^ → *m/z* 264.3 (282.3) for all Cer and [M+H]^+^ → *m/z* 184.1 (86.1) for all SM subspecies (C16, C18, C20, C22, C24 and C24:1) (114). Peak areas of Cer and SM subspecies, as determined with MassHunter Quantitative Analysis software (version 10.1, Agilent Technologies), were normalized to those of the internal standards (C17 Cer or d_31_-C16 SM) followed by external calibration in the range of 1 fmol to 50 pmol on column.

### Cytotoxicity Assays

For all cytotoxicity assays, HuLEC were seeded in 24 well plates with a density of 0.5×10^5^ cells per well two days prior to the experiment. Then, cells were infected with *S. aureus* JE2 with the indicated MOI as described in “*S. aureus* infection” and cytotoxicity was determined 21 h p.i. For cytotoxicity measurements upon ionomycin treatment, cells were incubated with the indicated concentration of ionomycin in infection medium for 75 min and then, cytotoxicity assays were conducted.

Lactate dehydrogenase (LDH) assay was performed with the Cytotoxicity Detection KitPLUS (LDH, Sigma Aldrich, Cat. No. 4744934001) according to manufacturer’s instructions.

For annexin V and 7-Aminoactinomycin D (7-AAD) assays, cells were washed thrice with DPBS, detached with 250 µL per well trypsin and resuspended in 250 µL per well staining buffer [1.7% (v/v) APC Annexin V (BD Pharmingen^TM^, Cat. No. 550475), 1.7% (v/v) 7-AAD (BD Pharmingen^TM^, Cat No. 559925), 2% (v/v) heat-inactivated (56°C/30 min) FBS (Sigma Aldrich, Cat. No. F7524), 4 mM CaCl_2_ (Roth, Cat. No. 5239.1) in DPBS]. After incubation for 10 min/RT, cells were analyzed with an Attune NxT flow cytometer for 7-AAD (Ex. 488 nm/Em. band pass 695/40 nm) and APC (Ex. 637 nm/Em. band pass 670/14 nm). Cell populations were analyzed with Flowing2 (Turku Biosciences Center) and gates were adjusted according to untreated control cells. To determine the proportion of cells that remained attached to the substratum during the infection, the number of cells was determined based on the number of detected single cells, flow rate and sample volume.

### Statistical analysis

Statistical analysis was performed in GraphPad prism (V10.1.2). One-sample t-test was used for analysis of normalized data sets. Otherwise, one- or two-way ANOVA, dependent on the number of variables, was used in combination with appropriate multiple comparisons testing. Details about sample size and corresponding statistical analysis can be found in respective figure legends. All data are shown as mean ± standard deviation.

## Supplementary Figures

**Supp. Figure 1.**
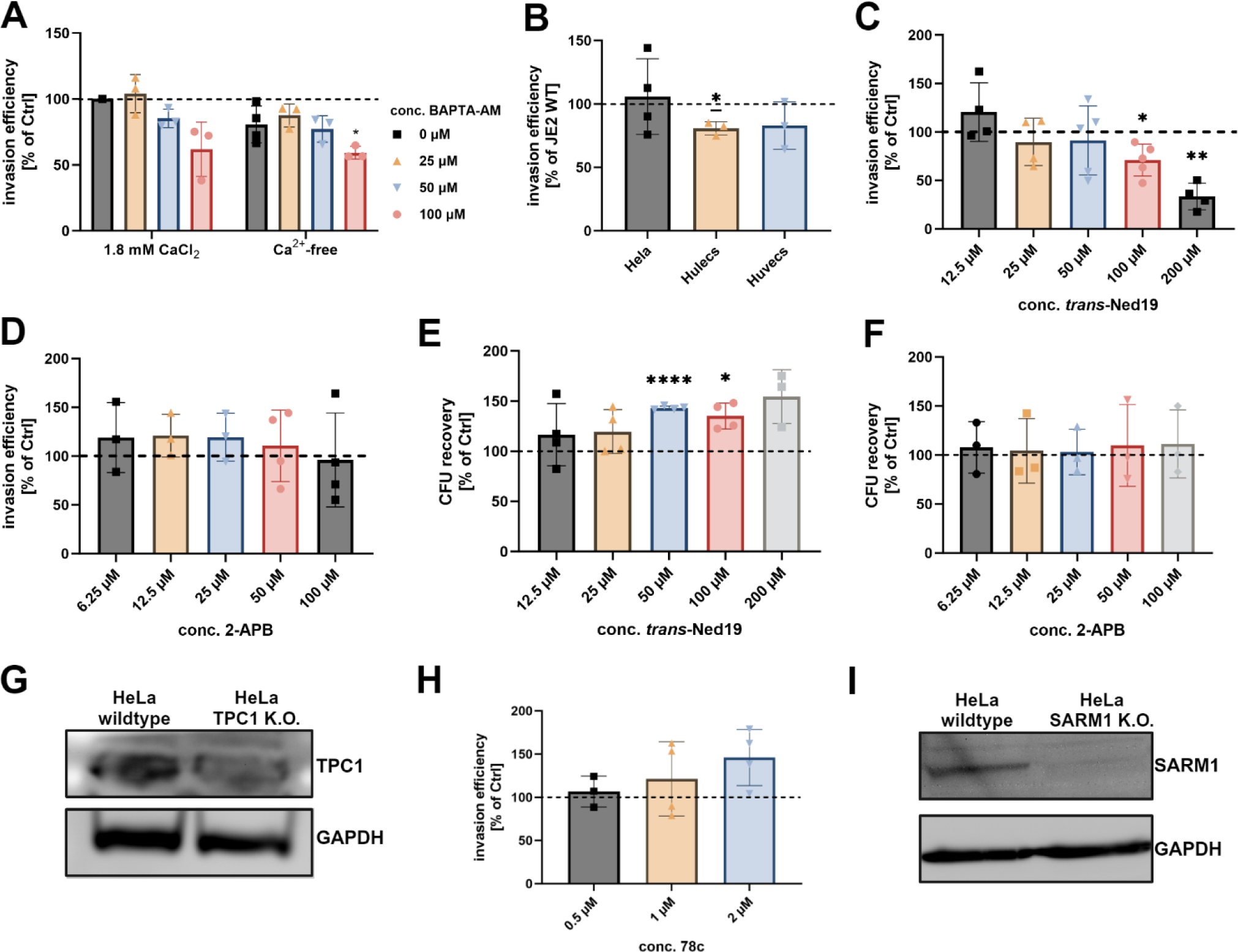
Supporting information for Ca^2+^-dependent internalization of *S. aureus*. (**A) Chelation of intracellular but not presence of extracellular Ca ^2+^ affects *S. aureus* internalization by HeLa cells.** HeLa cells were preloaded with varying concentrations of the cell-permeable Ca^2+^ chelator BAPTA-AM and then infected with *S. aureus* JE2 in presence (+1.8 mM CaCl_2_) or absence (Ca^2+^-free) of Ca^2+^ in the cell culture medium. Invasion efficiency was determined by CFU plating. (**B) α-toxin plays only a subordinate role during *S. aureus* uptake by host cells.** HeLa, HuLEC or HuVEC were infected with *S. aureus* JE2 wildtype or a strain deficient in α-toxin production (Δ*hla*). Invasion efficiency was determined by CFU plating. The number of intracellular bacteria was normalized to samples infected with the wildtype. **(C, D) *trans*-Ned19 but not 2-APB affects invasion of *S. aureus* JE2 in HeLa.** HeLa cells were pre-treated with *trans*-Ned19 (**C**) or 2-APB (**D**). 2-APB was removed from the cells shortly before infection to avoid direct contact of inhibitor and bacteria (due to a bactericidal effect of 2-APB; data not shown). Then, cells were infected with *S. aureus* JE2 and invasion efficiency was determined. **(E, F) *trans*-Ned19 and 2-APB have no effect on *S. aureus* JE2 in our infection protocol.** HeLa cells were pre-treated with or 2-APB (**E**) or *trans*-Ned19 (**F**). 2-APB was removed from the cells shortly before infection. Then, cells were infected with *S. aureus* JE2 and, after 30 min, total bacteria (extra- and intracellular) were recovered by CFU plating. (**G) Reduction of TPC1 expression in a Cas9-treated cell pool (HeLa TPC1 K.O.).** TPC1 as well as GAPDH (loading control) was detected in lysates of wildtype and TPC1 K.O. cells via Western blot. **(H) CD38 is not involved in *S. aureus* internalization.** HeLa cells were pretreated with the CD38 inhibitor 78c and invasion efficiency of *S. aureus* JE2 was determined. (**I) Reduction of SARM1 expression in a Cas9-treated cell pool (HeLa SARM1 K.O.).** TPC1 as well as GAPDH (loading control) was detected in lysates of wildtype and SARM1 K.O. cells via Western blot. Statistics: one sample t-test, Bars represent mean +/- SD. *p≤0.05, **p≤0.01, ***p≤0.001, ****p≤0.0001.

**Supp. Figure 2.**
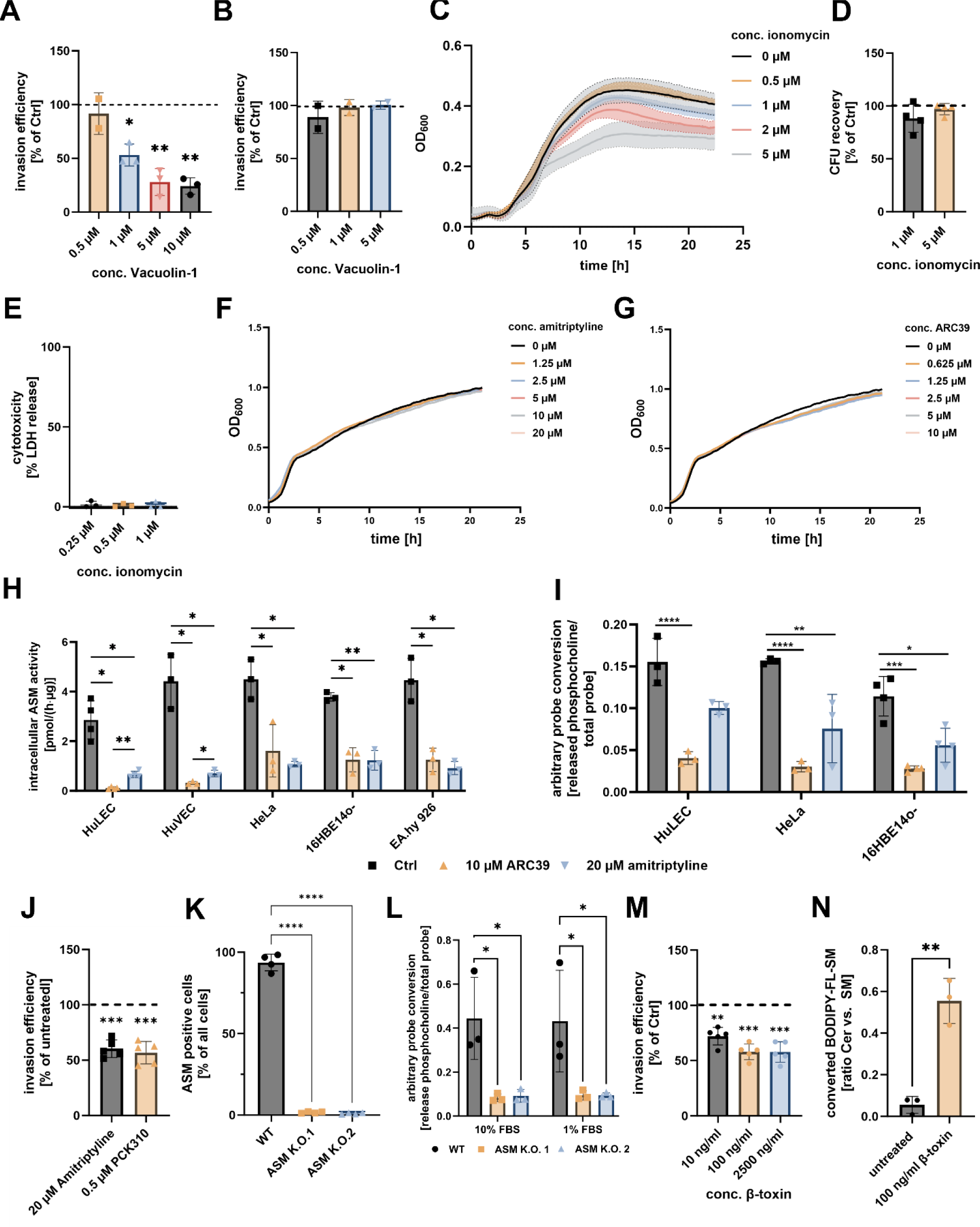
Supporting information for the involvement of lysosomal exocytosis and ASM in *S. aureus* invasion. **(A) High ionomycin concentrations interfere with growth of *S. aureus* JE2.** *S. aureus* JE2 was grown in in presence of varying concentrations of ionomycin. Growth (OD_600_) was determined in a microplate reader. (**B) Ionomycin does not affect survival of *S. aureus* JE2 during infection of host cells.** HeLa cells were pretreated with varying concentrations of ionomycin and subsequently, infected with *S. aureus* JE2. After 30 min, cells were lysed and surviving extra- and intracellular bacteria were recovered by CFU count. (**C) Ionomycin does not affect host cell survival.** HeLa cells were treated with the indicated concentrations of ionomycin, and cytotoxicity was determined by measuring in lactate dehydrogenase (LDH) release. **(D) Blocking lysosomal exocytosis by Vacuolin-1 reduces *S. aureus* invasion in HeLa.** HeLa cells were treated with increasing concentrations of the lysosomal exocytosis inhibitor Vacuolin-1. The invasion efficiency of *S. aureus* JE2 was determined 30 min p.i. **(E) Vacuolin-1 has no bactericidal effect.** HeLa cells were treated with Vacuolin-1 and infected with *S. aureus* JE2. After 30 min, total bacteria (extra- and intracellular) were recovered by CFU plating and normalized to untreated controls. **(F, G) The ASM inhibitors amitriptyline and ARC39 have no effect on growth of *S. aureus*.** *S. aureus* JE2 was grown in presence of varying concentrations of amitriptyline (**F**) or ARC39 (**G**) in BHI medium and growth was determined by measuring OD_600_ in a microplate reader. **(H, I) ASM activity and ASM inhibition by ARC39 and amitriptyline are similar among human cell lines.** Several cell lines were treated with amitriptyline or ARC39 and ASM activity within cell lysates was assessed by either thin layer chromatography-based ASM activity assays, detecting the conversion of BODIPY-C12-SM (**H**), or by a flow cytometry-based ASM activity assay and conversion of visible-range FRET probe (**I**). **(J) Microscopy-based measurement of invasion efficiency in amitriptyline- and PCK310-treated Hela cells.** Hela cells were treated with 20 µM amitriptyline (75 min) or 0.5 µM PCK310 (4h) and infected with *S. aureus* JE2 expressing Cerulean. Number of intracellular bacteria per host cell was determined by CLSM. n=5. **(K) Validation of ASM K.O. cell pools.** ASM activity in HeLa wildytpe or ASM K.O.s was determined with the visible-range FRET probe and flow cytometry. The proportion of ASM-positive cells in the cell population was determined. n=4. **(L) FBS concentration in culture medium does not affect cell-associated ASM activity in ASM K.O. cell lines.** HeLa WT or ASM K.O. cells were cultured in 1% or 10% FBS and cellular ASM activity was determined with a visible range FRET probe by flow cytometry. n=3. **(M) Removal of SM from the plasma membrane by β-toxin affects *S. aureus* host cell entry in HeLa.** HeLa cells were pretreated with the indicated concentrations of β-toxin and subsequently the invasion efficiency of *S. aureus* JE2 was determined. **(N) Treatment of HeLa with the bacterial SMases β-toxin results in generation of ceramide.** HeLa cells were preloaded with 1 µM BODIPY-FL-C_12_-SM and treated with β-toxin for 75 min. Quantities of ceramide and SM were determined by TLC and ratio of ceramide vs. SM were calculated. Statistics: one sample t-test (D, J, M), mixed-effects model (REML) and Tukey’s multiple comparison (H, I), one-way ANOVA and Dunnett’s multiple comparison (K), two-way ANOVA and Šídák’s multiple comparison (L), unpaired Student’s t-test (N). Bars represent mean +/- SD. *p≤0.05, **p≤0.01, ***p≤0.001, ****p≤0.0001.

**Supp. Figure 3.**
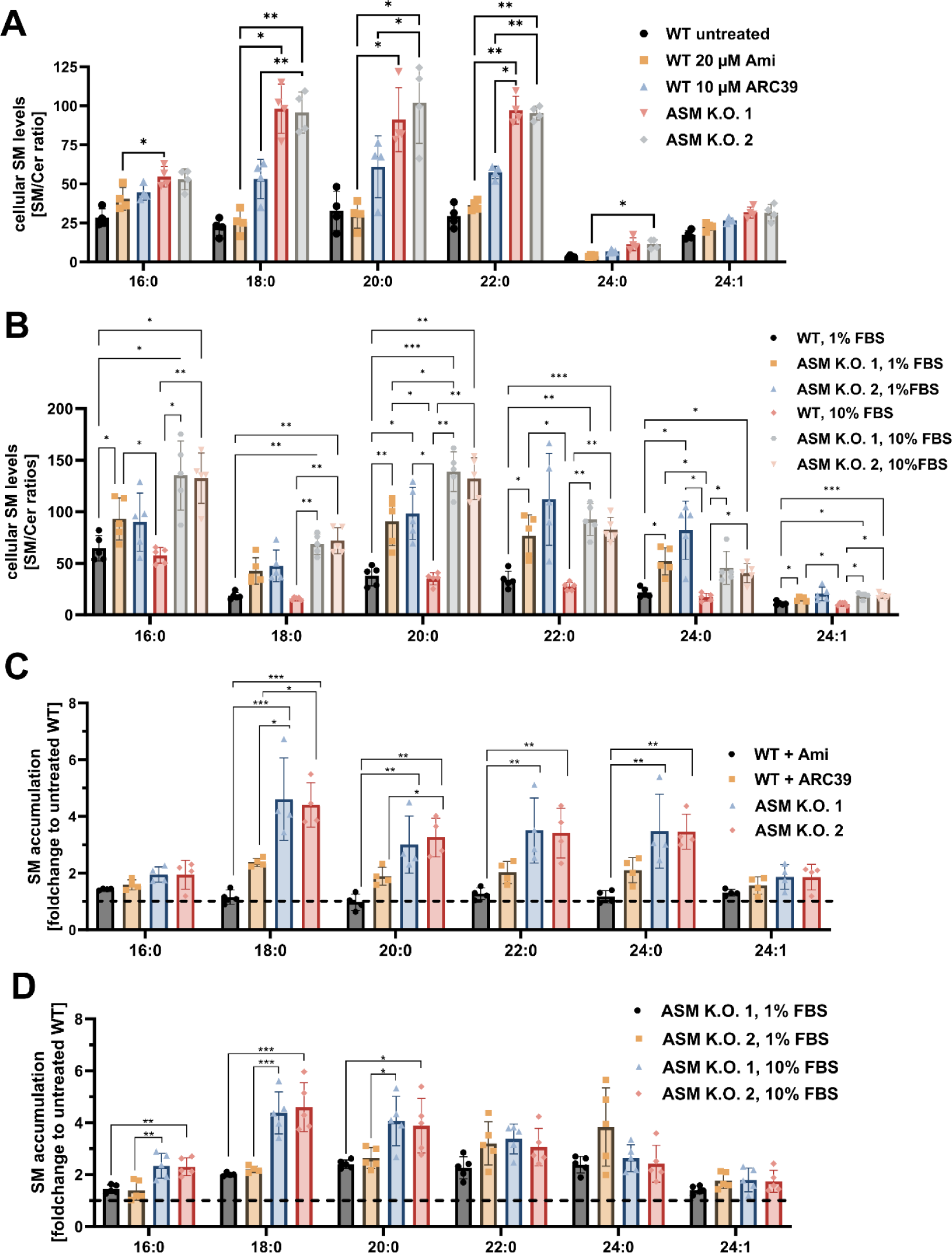
Effects of genetic K.O. and inhibition of ASM on cellular SM levels. HeLa WT or two ASM-depleted HeLa cells pools (ASM K.O. 1 and 2) were cultured in medium containing 10% or, if indicated, 1% FBS. If indicated WT cells were exposed to 20 µM amitriptyline for 75 min, or to 10 µM ARC39 for 22h. Amounts of SM and Cer species with different acyl chain lengths were quantified by HPLC-MS/MS and SM/Cer ratios were calculated to determine the cellular SM levels (A and B). To assess SM accumulation upon K.O. or inhibitor treatment, SM levels were normalized to untreated WT cells (set to 1, C and D). n≥4. Statistics: Two-way ANOVA and Tukey’s multiple comparison (A, B) and One-way ANOVA with Tukey’s multiple comparison for individual acyl chain lengths (C, D). Bars represent mean +/- SD. *p≤0.05, **p≤0.01, ***p≤0.001, ****p≤0.0001.

**Supp. Figure 4.**
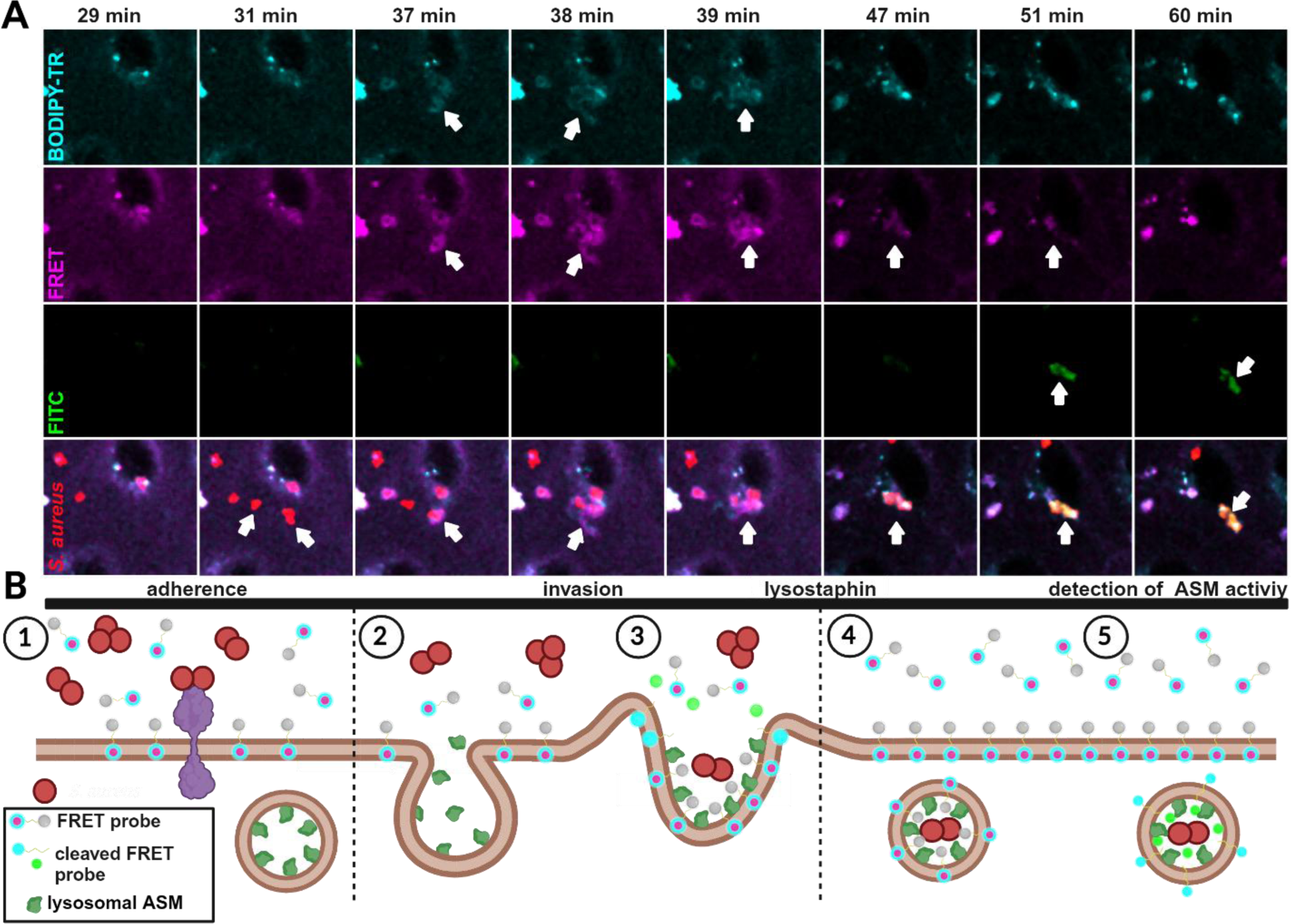
*S. aureus* associates with a visible-range SM probe during host cell invasion. **(A) Live cell imaging of *S. aureus* infection reveals association of the bacteria with a SM analog during cell entry.** HuLEC were treated with 10 µM of a visible range FRET probe in presence of 1% FBS. Then, cells were infected with *S. aureus* JE2 in presence of the probe and infection was monitored by live cell confocal imaging. After 40 min, lysostaphin was added to remove extracellular bacteria. The BODIPY-TR signal demonstrates the localization of the probe, the FRET signal indicates an intact, non-metabolized probe, whereas FITC fluorescence indicates probe cleavage by ASM. Bacteria of interest (white arrows) adhere to host cell between 29 and 31 min p.i. **(B) Hypothetical model of *S. aureus* invasion.** (1). The interaction of bacteria with the host cell surface triggers lysosomal exocytosis, release of ASM (2) and the uptake of the bacteria (cmp. to A: at 37-39 min p.i.) by ASM-dependent membrane remodeling (3). ASM is co-internalized together with bacteria (4) and subsequently, cleaves the probe within the *S. aureus*-containing phagosome (5) (cmp: to A: increasing FITC signal starting at 47 min p.i.).

**Supp. Figure 5.**
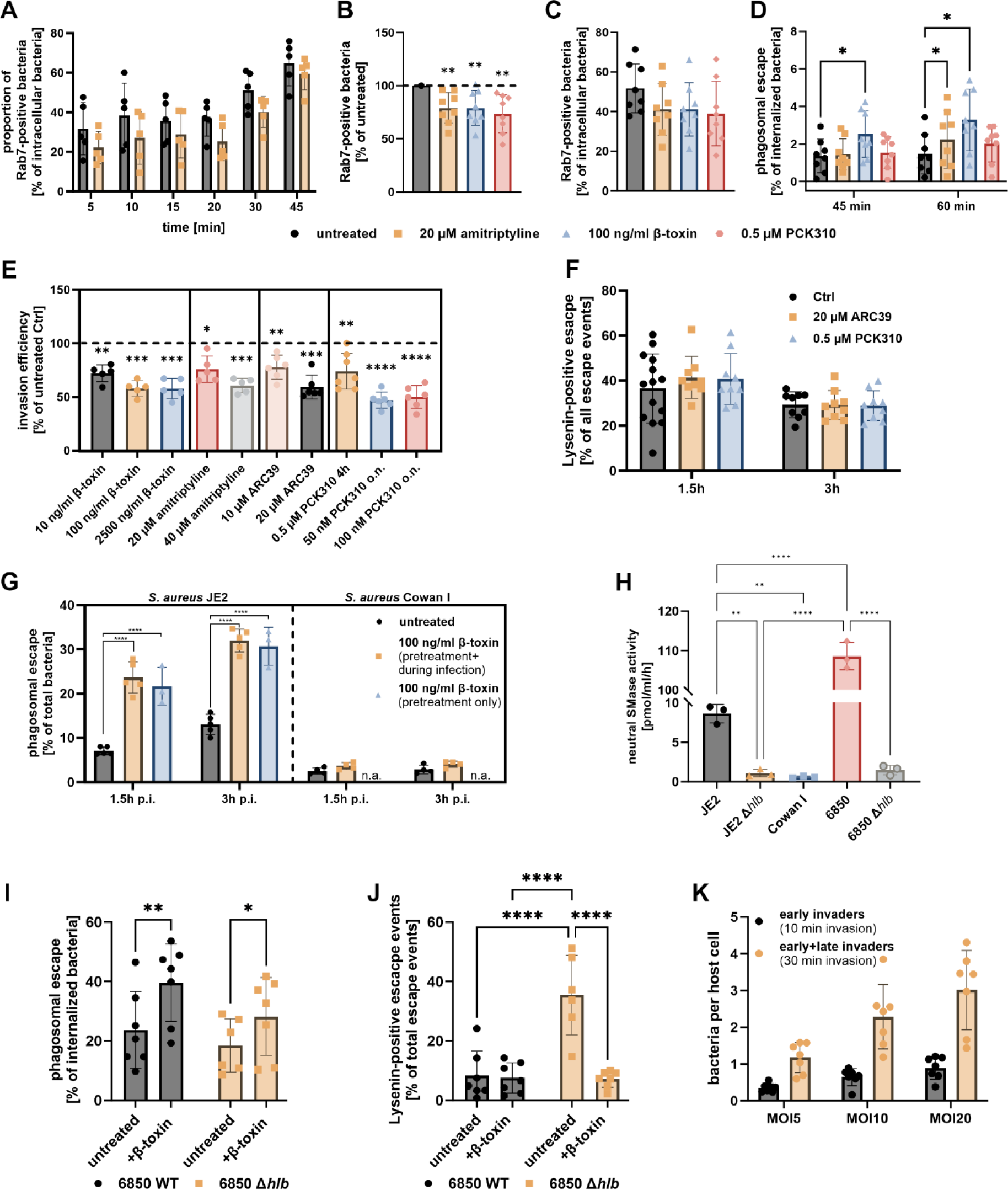
Supporting information for phagosomal maturation and phagosomal escape measurements. **(A) amitriptyline treatment reduces the proportion of Rab7-positive bacteria.** HeLa cells expressing YFP-Rab7 and mCherry-Rab5 were infected with *S. aureus* JE2 for the indicated time point and extracellular bacteria were removed with lysostaphin for 15 min. Then, samples were fixed and imaged by CLSM. Proportion of bacteria associated with YFP-Rab7 was determined. n=5. **(B, C) Reduced association of *S. aureus* with Rab7 is not due to changes in phagosomal escape.** HeLa cells expressing YFP-Rab7 and the phagosomal escape marker RFP-CWT were treated with amitriptyline, PCK310 or β-toxin and subsequently, infected with *S. aureus* JE2 for 45 min and analyzed by CLSMs. Bacteria that escaped from phagosomes were subtracted from the data set and the proportion of residual bacteria associated with Rab7 was determined (C). Results of treated samples were normalized to untreated controls (B, n=7). **(D) Phagosomal escape assay in early infection.** A HeLa reporter cell line expressing YFP-Rab7 and RFP-CWT was treated with amitriptyline, β-toxin and PCK310 and subsequently infected with *S. aureus* JE2 for 30 min. Extracellular bacteria were removed by lysostaphin and the proportion of bacteria that acquired the phagosomal escape marker RFP-CWT was determined 45 and 60 min p.i. (n=8). **(E) β-toxin treatment and ASM inhibition reduced invasion efficiency of *S. aureus* JE2 in a dual reporter HeLa cell line.** A HeLa reporter cell line expressing RFP-CWT and Lysenin^W20A^-YFP was treated with the indicated concentrations of β-toxin and amitriptyline for 75 min, the indicated concentrations of ARC39 for 22 h, 0.5 µM PCK310 for 4h or 50/100 nM PCK310 for 22h (o.n.) and invasion efficiency of *S. aureus* JE2 was determined. (n≥5) **(F) Inhibition of ASM has no influence on the proportion of Lysenin^W20A^-positive escape events.** The dual reporter cell line was infected with *S. aureus* JE2. Extracellular bacteria were removed and the proportion of bacteria that acquired the phagosomal escape marker RFP-CWT as well as the SM reporter Lysenin^W20A^-YFP was determined by CLSM. The proportion of escaped (RFP-CWT-positive) bacteria that additionally acquired Lysenin^W20A^-YFP was calculated. (n=9). **(G) β-toxin pretreatment increases phagosomal escape of *S. aureus* JE2 but not *S. aureus* Cowan I.** RFP-CWT and Lysenin^W20A^-YFP expressing HeLa were treated with 100 ng/ml β-toxin infected with *S. aureus* strains JE2 or Cowan I. Thereby, β-toxin was either removed prior to infection (pretreatment only) or was present during the whole experiment (pretreatment + during infection). By CLSM, proportions of bacteria that recruited RFP-CWT (phagosomal escape) were determined (n≥3). **(H) *S. aureus* strains vary in β-toxin expression.** The neutral SMase activity in culture supernatants of the indicated *S. aureus* strains or isogenic β-toxin mutants (Δ*hlb*) was determined by a TLC-based approach (n=3). **(I, J) β-toxin pretreatment of host cells but not β-toxin expression of endocytosed *S. aureus* affects phagosomal escape.** HeLa RFP-CWT/Lysenin^W20A^-YFP were infected with *S. aureus* 6850 or an isogenic strain deficient in β-toxin production (*Δhlb*). The proportion of bacteria that escaped from the phagosome (I) as well as the percentage of phagosomal escape events that were Lysenin^W20A^-positive (J) were determined 3h p.i. (n=6). **(K) Increasing the MOI compensates for lower numbers of invading bacteria at shorter infection times.** HeLa cells were either infected for 10 min or 30 min with varying MOIs. Then, the number of intracellular bacteria per host cell was determined 3h p.i. by CLSM. (n=7). Statistics: One samples t-test (B, E), mixed effects analysis (REML) and Dunnett’s multiple comparison (D), Two-way ANOVA with Tukey’s multiple comparison (G), One-way ANOVA with Tukey’s multiple comparison (H), mixed effects analysis (REML) and uncorrected Fisher’s LSD (I), mixed effects analysis (REML) with Šídák’s multiple comparison (J).Bars represent mean +/- SD. *p≤0.05, **p≤0.01, ***p≤0.001, ****p≤0.0001.

**Supp. Figure 6.**
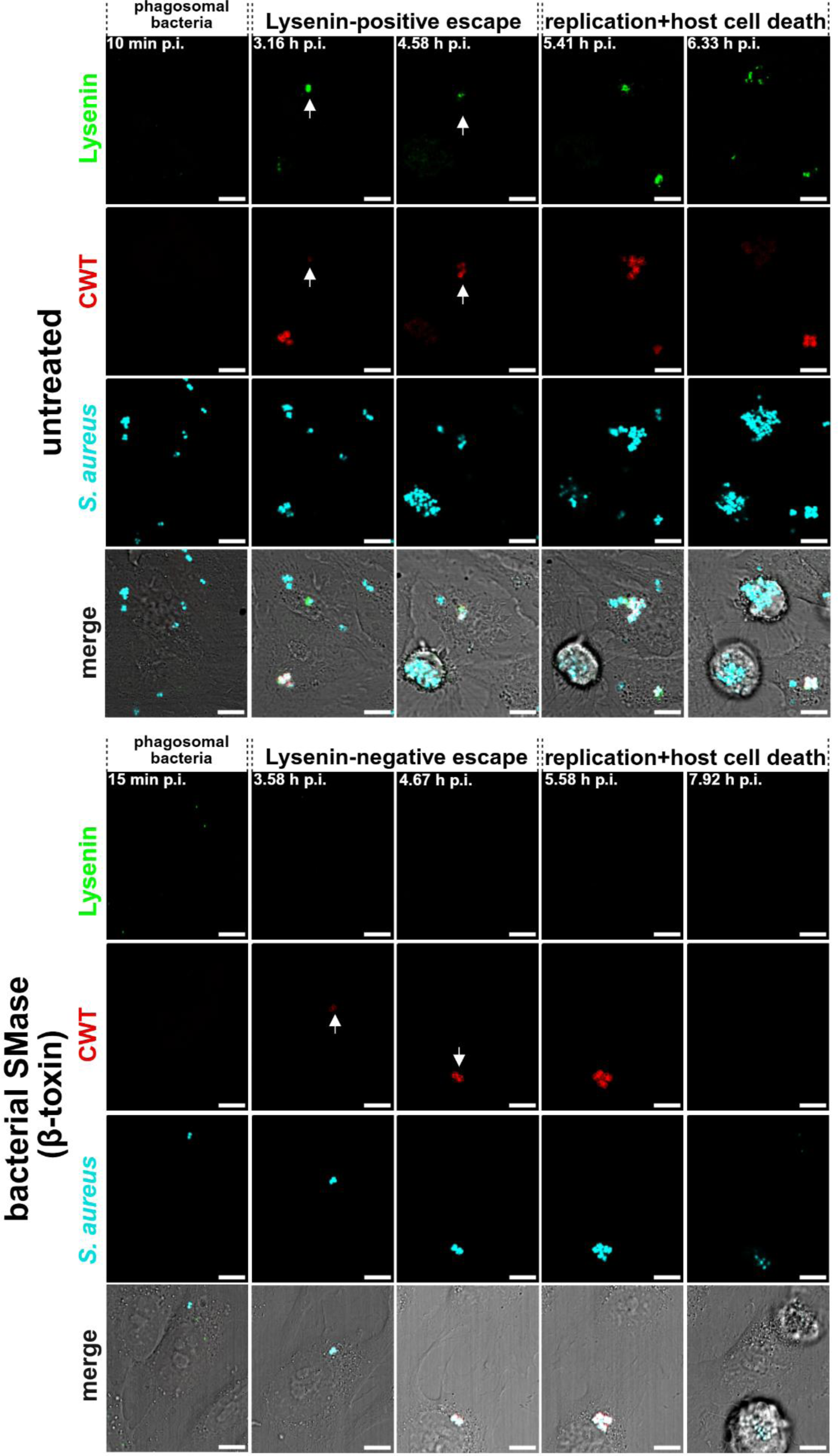
Validation of Lysenin^W20A^-YFP and RFP-CWT reporter cell line. Hela cells expressing the SM reporter Lysenin^W20A-YFP^(green) and the phagosomal escape reporter RFP-CWT (red) were either pretreated with 100 ng/ml bacterial SMase (A) or left untreated (B). Cells were infected with Cerulean-expressing *S. aureus* JE2 (cyan), extracellular bacteria were removed, and intracellular infection was monitored by time-lapse imaging at a Leica TCS SP5 microscope in 5 min intervals. White arrows indicate Lysenin-positive (A)or Lysenin-negative (B) escape events. Scale bars: 10 µm.

**Supp. Figure 7.**
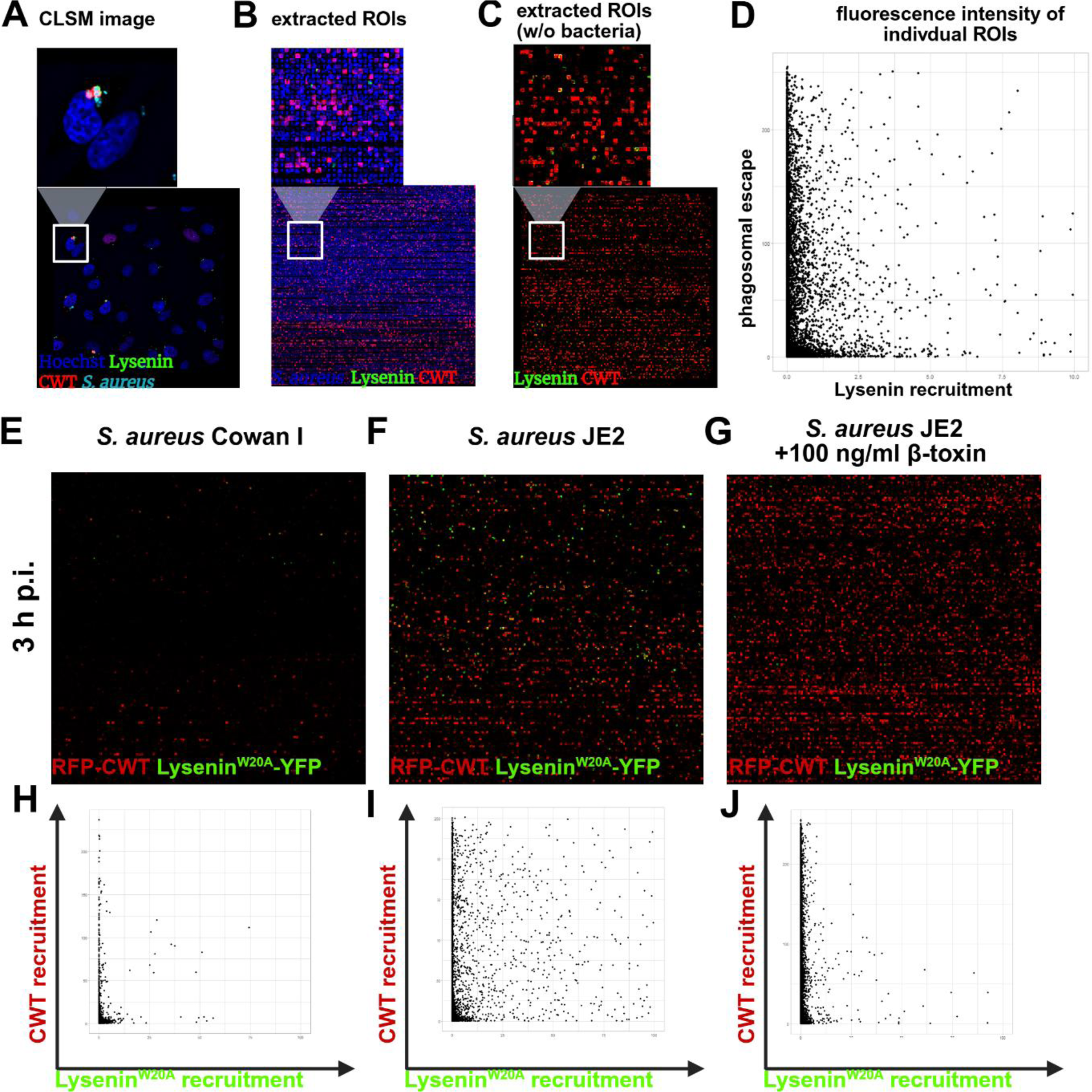
Evaluation of phagosomal escape and Lysenin^W20A^ recruitment during *S. aureus* infection. HeLa reporter cells expressing RFP-CWT and Lysenin^W20A^-YFP were infected with *S. aureus* JE2, fixed and imaged by CLSM (**A**). Individual bacteria within the images were identified as regions of interest (ROIs) and extracted in each color channel. All extracted ROIs are depicted in a montage (**B**). If bacteria signals are omitted, the montage of extracted ROIs shows the proportion of reporter recruitment. The average fluorescence intensity of both reporter fluorophores is measured within each individual ROIs is plotted as dot plots (**D**). From these results, the proportion of bacteria that recruited the markers (CWT-RFP = phagosomal escape and Lysenin^W20A^ = SM-rich-phagosome) is calculated. **(E-J) Exemplary escape assay.** Montages of extracted ROIs (without bacterial fluorescence) as well as the resulting intensity measurement in individual ROIs 3 h p.i. are depicted for infections of *S. aureus* Cowan I (E, H) and JE2 (F, I) as well as an infection with *S. aureus* JE2 where host cells were treated with 100 ng/ml β-toxin prior to infection (G, J). Data are quantified in **Figure 5, B, C**.

**Supp. Figure 8.**
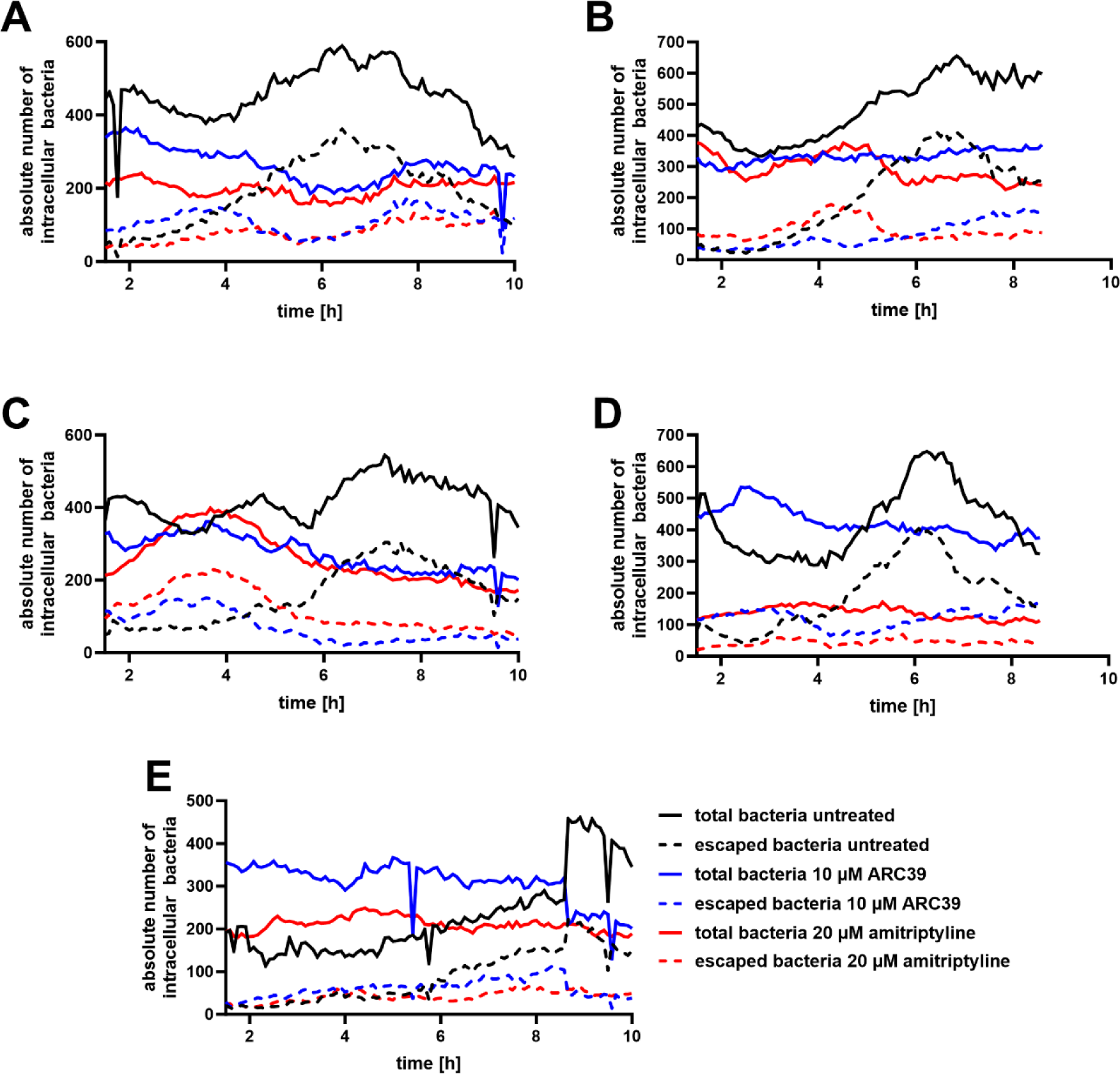
Individual replicates of time-dependent phagosomal escape and replication assays. HeLa cells expressing RFP-CWT were pretreated with 20 µM amitriptyline or 10 µM ARC39, infected with *S. aureus* JE2 and infection was monitored by live cell imaging (for further details see **Figure 6, A** and B). The absolute numbers of all intracellular bacteria and bacteria that escaped from the phagosome, which were detected during live cell imaging, are depicted.

**Supp. Figure 9.**
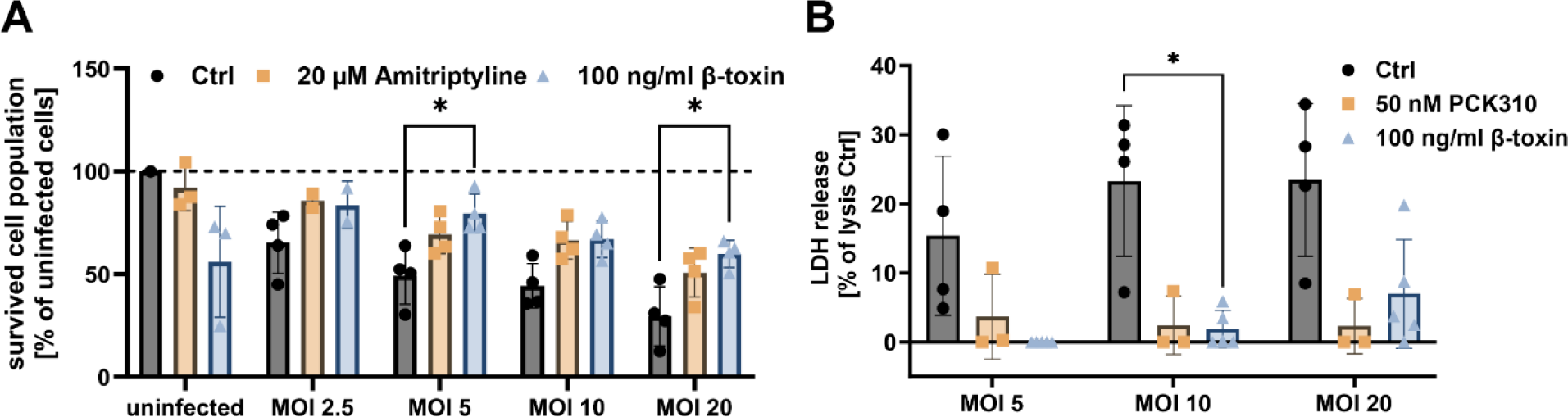
Inhibition of ASM and β-toxin treatment increase number of host cells remaining attached to the substratum and reduce host cell lysis during infection. HuLEC were treated with amitriptyline or β-toxin (**A**) or PCK310 and β-toxin (**B**) and were then infected with *S. aureus* JE2. After 30 min, extracellular bacteria were removed and the number of cells remaining attached to the substratum was determined (**A**) or the proportion of host cells that were lysed during the infection was measured by lactate dehydrogenase (LDH) release (**B**). Statistics: Mixed-effects analysis and Dunnett’s multiple comparison (A, B). Bars represent mean +/- SD. *p≤0.05, **p≤0.01, ***p≤0.001, ****p≤0.0001.

## Supplementary Videos

**Supp. Video 1 *S. aureus* associates with BODIPy-FL-C_12_-sphingomyelin during invasion** (for further details, see Figure 3, A)

**Supp. Video 2 *S. aureus* associates with a visible-range FRET probe during invasion** (for further details, see Supp. Figure 4)

**Supp. Video 3 *S. aureus* associates with Rab5 and Rab7 upon host cell entry** Hela cells expressing mCherry-Rab5 (red) and YFP-Rab7 (green) were infected with fluorescent *S. aureus* JE2 (cyan). Time-laps imaging was performed at a Leica TCS SP5 microscope in intervals of 45s. Scale bar: 10 µm.

**Supp. Video 4 Validation of Lysenin and CWT reporter cell line** (for further details see Supp. Figure 6)

**Supp. Video 5 β-toxin pretreatment protects HuLEC from *S. aureus* infection** (for further details see Figure 6, c)

